# Plant phenotypic plasticity changes pollinator-mediated selection

**DOI:** 10.1101/2022.02.10.479857

**Authors:** Thomas Dorey, Florian P. Schiestl

## Abstract

Many organisms change their phenotype in response to the environment, a phenomenon called phenotypic plasticity. Yet we hardly understand how such plasticity can affect biotic interactions and the resulting phenotypic selection. Here we use fast cycling *Brassica rapa* plants in a proof-of-concept experiment in the greenhouse to study the link between plasticity and selection. We detected strong plasticity in morphology, nectar, and floral scent in response to different soil types and aphid herbivory. We found positive selection on nectar and several morphological traits. Bumblebee-mediated selection on a principle component representing plant height, flower number, and flowering time (mPC3) differed depending on soil type and herbivory. For plants growing in richer soil, selection was stronger in the absence of herbivores, whereas for plants growing in poorer soil selection was stronger with herbivory. We showed that bumblebees visited tall plants with many flowers over-proportionally when they were rare (i.e. in plants in poor soil with herbivory), thus causing stronger positive selection on this trait-combination. We suggest that with strong plasticity under most stressful conditions, a shift in pollinator behavior may speed up adaptation to local environmental factors.

## Introduction

Environmentally-induced change in an organisms’ phenotype, so-called phenotypic plasticity, is commonly found among all kinds of organisms (West-Eberhard 1989, Agrawal 2001, Pfennig 2021). In plants, plasticity is known to be induced by abiotic-or biotic factors, affects many different kinds of traits, and is commonly classified as being adaptive or non-adaptive (van Kleunen and Fischer 2005). Abiotic stress can induce changes in floral display (Phillips et al. 2018), flower-reward (Descamps et al. 2021), nectar- and leaf chemistry (Mattson and Haack 1987, Hoover et al. 2012), flower development (Hoover et al. 2012), plant biomass as well as plant volatile emission (Luizzi et al. 2021). For biotic factors, herbivory is one of the most commonly studied factors triggering phenotypic plasticity in plants, typically leading to increased direct defense, such as toxicity or trichome density (Agrawal 1999b), or indirect defense through attraction of the herbivores’ enemies (Turlings et al. 1990, Kessler and Baldwin 2001).

Because plants interact through their phenotype with other organisms, phenotypic plasticity may impact such interactions in various ways. For example, abiotic stress can impose changes on plant-pollinator (Harrison 2000, Burkle and Runyon 2016) as well as plant-herbivore interactions (Mattson and Haack 1987, Price 1991). Herbivore-induced defenses are known to alter pollinator behavior, e.g. through changing flower signals and/or nectar chemistry, usually rendering flowers less attractive to pollinators (Lehtila and Strauss 1997, Adler et al. 2006, Kessler et al. 2011, Schiestl et al. 2014, Rusman et al. 2020), but see (Poveda et al. 2003, Cozzolino et al. 2015, Goulnik et al. 2020). Other documented effects of herbivore-induced plasticity are shorter visitation times of pollinators (Bruinsma et al. 2014, Ramos and Schiestl 2019), changes in pollination networks (Hoffmeister et al. 2016), and even switches in pollinator guilds attracted to flowers (Kessler et al. 2010).

Although environmental factors such as soil or herbivory can directly impose selection on plants, for example through limiting nutrients for seed production or direct destruction of seeds or ovules (Strauss et al. 2002), modifying key interactions such as pollination is a potentially powerful indirect way of impacting natural selection in plants. Among the various biotic interactors, pollinators are a strong selective agent through their specific morphology, sensory abilities, and behavior, that directly impact the plants’ reproductive success (Strauss and Irwin 2004, Schiestl and Johnson 2013, Sletvold et al. 2017, Caruso et al. 2019). Although selection/adaptation studies often circumvent the effects of pollinators by enabling selfing model plants (Mitchell-Olds and Schmitt 2006, Agren et al. 2017), the majority of angiosperms requires animal pollinators for sexual reproduction to variable degrees (Ollerton et al. 2011), and thus pollinator-mediated selection will contribute somehow to their adaptive evolution. How biotic and abiotic factors impact pollinator-mediated selection is, however, hardly understood (Herrera 2000, Gomez 2003, Caruso et al. 2005) and thus we know little about how environmentally induced plasticity can impact adaptive evolution indirectly, via its modifying effect on selection.

In this study, we investigate the importance of biotic and abiotic agents on plant phenotypic plasticity, and its subsequent effect on phenotypic selection. We first quantify the plants’ responses to two different soil types and the presence/absence of aphid herbivory in a full factorial design. Then, by using phenotypic selection analyses, we evaluate how the soil- and herbivory-induced changes affect phenotypic selection imposed by bumblebee pollinators. Our approach aimed to determine how three different selective agents (soil, herbivore, pollinator) interact in shaping natural selection and thus potential plant adaptative evolution. We addressed the following specific questions: (1) How do plants respond to different soil types and to herbivory in terms of phenotypic plasticity? (2) Does phenotypic plasticity alter pollinator-mediated selection on plant traits?

## Material and Methods

### Study system

We used outcrossing, annual “fast cycling” *Brassica rapa* plants (Wisconsin Fast Plants® Standard Seed with maximum genetic diversity) as a model system for our study. These plants have a short generation time, comprise high genetic variation, are self-incompatible, and have been successfully used for selection and experimental evolution studies (Tomkins and Williams 1990, Gervasi and Schiestl 2017, Schiestl et al. 2018, Ramos and Schiestl 2019). To generate the plants used in this study, in 2018, we obtained 440 seeds from Carolina Biological Supply (Burlington, NC, USA) and grew them in a phytotron under standardized light, soil, temperature, humidity, and watering conditions. Of these 440 seeds, 410 germinated and were used to generate full sib seeds families by manually crossing 205 randomly assigned pairs. From these crossings we obtained a total of 163 full sib seed families (only pairs where both parents produced fruits were used in the experiment). Among those 163 seed families, we randomly chose 98 families to represent the starting population of our experiment.

This study was done within the first two plant generations of longer lasting experimental evolution study (Dorey and Schiestl, in prep); thus, all plants used for this experiment were grown in two subsequent generations. For the first generation, we assigned randomly the 98 families to two replicates (A and B) and 8 treatments (see below) so that each replicate included 49 plants (7×7, to allow rectangular setup in bee cage) representing 49 families (one plant per family). In each treatment, replicate A represented families (1-49) and replicate B represented families (50-98), so that the families were equally spread across the whole experiment. The experiment consisted of 8 treatments ã 2 replicates, and a total of 784 plants per generation. The second generation of plants was generated from the seeds produced by plants of the first generation through bumblebee pollination, where each individual contributed proportional to its total seed set, to achieve a total sample size of again 49 plants per replicate. Thus, generation two represented the response to selection in generation one. We incorporated the effect of this change in genotypic composition on selection in our statistical analyses, where the factor “generation” was not significant, suggesting that selection is consistent even with a change of genotype between generations (see results part for details). For both generations, plants were first grown in a phytotron under 24 hours of light, 21°C, 60% humidity and were watered once day (at 8:00). After pricking, the plants were transferred to an air-conditioned greenhouse with natural and artificial illumination to achieve 16h of light, at a constant temperature of 23°C.

### Selection experiment and treatments

Plants were grown either in limestone or tuff soil (Table S1). Both soils are found in different regions of Southern Italy, with native a species of *Brassica, B. incana*, growing in both of them. Soil was collected in July 2018 and October 2018 at two locations in Campania (Southern Italy); limestone: Valico di Chiunze: 40.719°N, 14.619°E, and tuff: Monte di Procida :40.809°N, 14.045°E. The locations were chosen because they represented two typical soil types of the region, and supported naturally growing *Brassica incana* plants, which were also investigated in another study (Arrigo et al in prep.). Soil (ca. 500 kg) was collected from the surface layer (depth 0-15cm), filtered using a mesh of 1cm of diameter, stored in bags and shipped to Switzerland. A sample of both soils was analyzed by the INRA laboratories following the SOL-1031 protocol (Table S1)(Ciesielski et al. 1997). This method used a solution of HF-HLCO4, an acid mixed which allowed to digest soil clay.

As the second treatment factor, in each type of soil (L: limestone; T: tuff), the plants were either exposed to herbivory by aphids (H: with herbivory) or kept without aphids (NH: no herbivory). Thus, our four different treatments (LNH, LH, TNH, TH) represented the factors soil and aphid herbivory in full factorial way. Aphids are the most common herbivores found in natural *Brassica incana* populations in Southern Italy (Arrigo and Schiestl, unpublished data). As aphid herbivore we used the *Brassicaceae’s* specialist, *Brevicoryne brassicae*. Aphids were collected in summer at the Botanical Garden of the University of Zurich (Switzerland) and then reared in climatic chambers on *Brassica rapa* under 16h of light at a temperature of 23°C and humidity of 70%. The infestation of the plants was done at the two true leaves stage which is about 13-14 days after sowing out for plants growing in tuff soil, and about 17-18 days in limestone. Ten wingless aphids were manually put on each plant and allowed to feed on the plants for 72 hours. During those 72 hours all plants (with and without aphids) were covered by a net (7×7×15cm) with a mesh of 680 µm (Bugdorm, model DC0901-W). After this period, aphids were manually removed from the plants and plants were subsequently checked every day for aphids that had accidently been left on the plants.

For pollination we used either random hand pollination or the bumblebee *Bombus terrestris*, a common generalist pollinator of *Brassica rapa* (Zu and Schiestl 2016, Ramos and Schiestl 2019). For hand-pollination, half of the plants per replicate were randomly chosen for pollinated, to achieve a similar proportion of plants with fruit/seed set as in the bee-pollinated treatments. Pairs of parent plants were randomly assigned; of four flowers per plant, one long stamen was removed and an excess pollen deposited on one stigma of a total of four flowers per each mother plant, so that each mother plant was pollinated with one father plant. Bumblebee hives were purchased in Switzerland (Andermatt Biocontrol Suisse AG). The hives were kept in a flight cage (75×75×115cm) and the bumblebees were allowed to feed on *Brassica rapa* fast cycling flowers, with supplemental pollen given (Biorex, Ebnat-Kappel, Switzerland) and additional solution of sugar water (Biogluc sugar solution, Biobest). When they arrived, bumblebees did not have any experience with plants. Therefore, we fed them for a week with flowering *Brassica rapa* plants to allow them to gain experience with these flowers. Plants were removed three days before pollination and bumblebees where from this moment only fed with sugar solution and additional pollen. To enhance the foraging activity of the bumblebees during pollination, we starved them 16 hours before pollination.

Pollination occurred 10 days after herbivory was terminated. For each replicate, pollination started at 8.30am and ended at 5.30pm and was set up under artificial light conditions. Plants were randomly placed in a square of 7×7 plants in a flight cage (l × w × h: 2.5m × 1.8m × 1.2 m) so that every plant was 20 cm apart from each other. A total of seven bumblebees were then allowed to subsequently forage on the plants. Bumblebee workers were released subsequently and recaptured after visiting a maximum of 5 different plants per individual; each insect was used only once. The number of visits per bee was limited to assure pollen limitation of reproductive success at the replicate level, pollen limitation being a common phenomenon among angiosperms in nature (Larson and Barrett 2000, Ashman et al. 2004). At the end of the pollination, all fully open flowers in visited plants were marked and only fruits that developed from these marked flowers were collected and counted after ripening.

After pollination, all visited plants were kept in the greenhouse under standardized light/watering conditions for completing fruit development. After four weeks the plants were then deprived of water and started to dry. Fruits and seeds harvested from the different pollination treatments were counted and total seeds weight measured.

### Resource limitation of seed production

To determine if soil nutrients were a limiting factor in reproductive success, we analyzed hand-pollinated without herbivory. We also regrew 49 plants (one individual of all full sib families) in both soil types, with adding nutrients in the form of fertilizer, and compared seed set of plants with and without fertilizer after hand pollination (as described above). Nutrient addition consisted of 10mL of an universal garden fertilizer (NPK: 8-6-6 with traces of B, Cu, Fe, Mn, Mo, Zn; Wuxal, Maag Agro, Dielsdorf, Switzerland) diluted in 10L of water. The data showed a significant increase of seeds per silique from 8.57± 4.34 to 11.81 ± 4.72 (t-ratio: 3.809, p-value=0.001) in limestone soil, whereas in tuff soil the increase was non-significant from 10.27 ± 4.35 to 12.52 ± 5.51 (t-ratio: 2.577, p-value=0.051; Figure S1). Thus, we assume that seed production was resource limited only for plants growing in limestone soil in our experiment (we grew all plants for the selection experiment without fertilizer).

### Measuring plant traits

All plants traits (morphology and scent) were measured within five days before the pollination day. Scent collection was done first, to avoid altering the scent profile of the plants because of tissue injuries. As our morphological measurements were done in a destructive way (three flowers were picked), we stopped all morphological measurements three days before pollination to allow new flowers to open. After the pollination of a replicate was done, we phenotyped those plants which did not get pollinated and started flowering only within the three days before pollination, on the same day.

For measuring plant floral morphology (petal display: petal width, petal length, petal area, flower diameter and UV-reflecting area; reproductive organs: long stamen length, pistil length), we sampled three flowers per plant; we then placed petals and reproductive parts on a white sheet and took a picture with a UV-light sensitive digital camera (Nikon D200 digital SLR, with quartz lens: Nikon UV Nikkor f4.5/105mm with Baader 2” U-Filter (310-390nm UV transmission). Flowers were illuminated with a UV torch (1W-UV Torch, λ= 365 nm and intensity= 6.6 mW/cm2). Floral traits including UV-reflecting area were quantified from the pictures using the software package Image J (https://imagej.nih.gov/ij/). For quantitative analyses, the mean values, calculated from three sampled flowers, were used. Floral nectar was collected from the same three flowers with a 1 μL microcapillary (Blaubrand, Wertheim, Germany) which allowed assessing of nectar volume. Total nectar volume was then divided by number of sampled flowers for obtaining the mean nectar amount per flower.

Height and number of open flowers were measured/counted on the pollination day. For comparing the plants in the two soil types, we also measured the height of plants grown in limestone at the same age as the plant grown in tuff soil (because plants at the pollination day were not of the same age, because the delayed development in limestone soil). However, these two height measures were highly correlated (R^2^= 0.92) so only height at pollination day was kept for further analyses.

Plant scent was collected in a non-destructive way (Schiestl et al. 2014, Ramos and Schiestl 2019) from the whole plants’ inflorescence as soon as the plants had at least 2 open flowers, using headspace sorption. The inflorescence was enclosed in a glass cylinder previously coated with sigmacote (Sigma-Aldrich, Buchs, Switzerland) and closed with a magnetic Teflon locker. The lock-mechanism had in its center a hole of 0.5 cm diameter allowing the stem of the plant to fill the space without being injured. Air from the surrounding was pushed through an activated charcoal filter into the glass cylinder with a flow rate of 150 ml/min, to purify the incoming air. Simultaneously, air was pulled off the cylinder at a flow rate of 150 ml/min through a glass tube filled with 30 mg Tenax TA (60/80 mesh; Supleco, Bellefonte, PA, USA) to adsorb volatiles. Volatiles were collected for two hours in the greenhouse under controlled light and temperature. Air from empty glass cylinders was also collected in the same ways as air controls. During the process, flower numbers within the glass tube was counted to later standardize the volatiles amount emitted per flower and per unit of collected air volume. Volatiles samples were analyzed by gas chromatography with mass selective detection (GC-MSD; see below). Samples were inserted into a GC (Agilent 6890 N; Agilent Technologies, Santa Clara, CA, USA) by a MultiPurpose Sampler (MPS; Gerstel, Müllheim, Germany) using Gerstel thermal desorption unit (TDU, Mühlheim, Germany) and a cold injection system (CIS; Gerstel). The GC was equipped with a HP-5 column (15 m length, 0.25 mm ID, 0.25 µm film thickness) and used helium as purge gas at a flow rate of 2 ml min−1. Volatiles organic compounds were extracted from samples by thermodesorption. During this process, the TDU was heated from 30 to 240°C at a rate of 60°C min−1 and maintained at the final temperature for 5 min. The CSI was then set to −150°C while eluting compounds from the TDU were trapped. For injection, the CIS was heated to 250°C at a rate of 12°Cs-1 and maintained final temperature for 3 min. A mass selective detector (Agilent MSD 5975) was used for compound identification and quantification. Each compound was identified by comparing its mass spectra with those of the National Institute of Standards and Technology (NIST) mass spectral library. The retention time of the spectra and its amplitude were two main features allowing to compare our samples to those of the library. Three different concentration (1, 10 and 100 ng/L) of synthetic standards of all compounds were previously analyzed on the GC-MSD system to establish a quantification database. The quantification of the different compounds of our samples was done by using a calibration curve for target ions specific to the individual scent compounds. This way we were able to calculate the amount of each compound in our sample. Only compounds that were significantly different between air control and plant samples were included in the subsequent statistical analysis. All amounts of volatiles were calculated in pg flower^−1^ l sampled air^-1^.

### Statistical analysis

In order to approach normal distribution and homogeneity of variances, nectar and scent variables were logarithmically transformed using the formula log(1+x); variables of morphological traits were not transformed. To simplify our data set and to obtain orthogonal (uncorrelated) variables, we ran principal components analyses (PCA) on all traits. Initial PCA showed that morphological and scent variables were associated to different axes and could by consequence be studied separately (Table S2). Therefore, we ran principal component analyses on morphological and scent variables separately. To facilitate the interpretation of the obtained principle components (PCs), we performed rotations of the axis by using the varimax method to preserve the uncorrelatedness of the transformed component, yet maximize factor loading thus easing the interpretation of PCs. This analysis resulted in four principle components (PCs) with an eigenvalue above 0.9 for both scent and morphological profiles, explaining 72.1 and 71.6 % of the total variation, respectively (Table 1).

**Table 1.** Rotated loadings of the morphological and scent variables on the principal components of Brassica rapa in generation one and two combined. The highest loadings of each variable are indicated in bold. Only principal components with eigenvalues> 0.9 and that explain> 4 % of the total variance were retained

| Principal components for morphology |  |  |  |  | Principal components for scent |  |  |  |  |
| --- | --- | --- | --- | --- | --- | --- | --- | --- | --- |
|  | mPC1 | mPC2 | mPC3 | mPC4 |  | sPC1 | sPC2 | sPC3 | sPC4 |
| Eigenvalues | 4.65 | 1.78 | 1.29 | 1.20 | Eigenvalues | 5.87 | 1.4 | 0.99 | 0.94 |
| Variance explained (%) | 38.74 | 14.86 | 10.8 | 10.05 | Variance explained (%) | 45.16 | 10.78 | 7.63 | 7.20 |
| Rotated loading on retained components |  |  |  |  |  |  |  |  |  |
|  | PC1 | PC2 | PC3 | PC4 |  | PC1 | PC2 | PC3 | PC4 |
| Plant height | 0.19 | -0.03 | **0.82** | 0.23 | Benzaldehyde | **0.67** | 0.32 | 0.30 | -0.05 |
| Flowering time | -0.15 | -0.19 | **-0.76** | 0.10 | 1- Butene-4-<br>isothiocyanate | -0.01 | **0.84** | -0.04 | 0.12 |
| No. of open<br>flowers | 0.05 | -0.20 | **0.73** | 0.15 | Methyl benzoate | **0.79** | 0.06 | 0.05 | 0.31 |
| Nectar amount<br>(μL) | 0.05 | 0.06 | 0.06 | **0.77** | Phenylethyl alcohol | 0.23 | 0.20 | **0.82** | 0.24 |
| Pistil | 0.20 | **0.83** | 0.00 | 0.33 | 2-Amino<br>benzaldehyde | 0.34 | -0.15 | 0.45 | **0.66** |
| Stamen | 0.20 | -0.07 | 0.09 | **0.77** | p-Anisaldehyde | **0.62** | 0.10 | 0.29 | 0.20 |
| Anther- stigma<br>distance | 0.02 | **0.92** | -0.04 | -0.13 | Methyl anthranilate | 0.57 | -0.13 | 0.26 | **0.59** |
| Petal width | **0.75** | 0.02 | 0.20 | 0.31 | (Z) 3-Hexen-1-ol.<br>acetate | 0.28 | **0.63** | 0.19 | 0.00 |
| Petal length | **0.60** | 0.25 | 0.15 | **0.60** | Phenylacetaldehyde | 0.19 | 0.03 | **0.86** | 0.26 |
| Petal area | **0.93** | 0.07 | 0.15 | 0.15 | Benzyl nitrile | 0.35 | -0.05 | 0.50 | **0.66** |
| UV reflecting<br>area | **0.91** | 0.06 | 0.08 | -0.01 | Methyl salicylate | **0.71** | 0.11 | 0.12 | 0.37 |
| Flower<br>diameter | **0.60** | 0.21 | 0.13 | 0.56 | Indole | 0.19 | 0.08 | 0.31 | 0.74 |
|  |  |  |  |  | (E,E)-α-Farnesene | 0.15 | 0.35 | 0.01 | **0.77** |

To analyze visitation and seed set of plants in association with values of mPC3 in the limestone treatment, mPC3 was transformed into 5 categories for both limestone treatments (i.e. with- and without herbivory). The 5 categories (1: smallest plants, fewest flowers, 5: tallest plants, most flowers) contained almost equal overall number of plants: 1: 62; 2-4: 63, 5: 61, but the two treatments differed in the number of plants per category (see Figure 4 legend for numbers of plants in all categories).

### Phenotypic plasticity

In order to analyze phenotypic plasticity, we studied how traits varied between treatments by using the rotated principal components (PCs) scores in plants of generation one only. We used linear mixed models with traits as dependent variable, traits as covariates, treatment as fixed factor and replicate as random factor. Phenotypic plasticity at the individual trait (rather than PC) level was also tested and results are provided in supplementary Table S4.

We also graphically plotted the plasticity responses (reaction norms) of the different full-sib seed families to our treatments (Figure S2 and S3). Using the rotated components score (PCs), we represented only families where we could link specific treatments, namely TNH to TH, and/or TH to LNH and/or LNH to LH. In the graphs, a decrease in phenotypic values is represented by a red line whereas a blue line describes a reduction in phenotypic values among individuals of the same full sib seed family.

### Phenotypic selection

To analyze phenotypic selection, we analyzed associations between female fitness and principal component (PC) scores in plants of generation one and two combined. For doing so, we used linear general mixed models with relative seed set (as fitness component) as dependent variable, traits as covariates, “soil”, “herbivory”, and “pollination” (the latter was included only for the full dataset with hand- and bee-pollination) as fixed factor and replicate as random factor. Generation was first included in all models as fixed factor; because generation was not significant we omitted it in subsequent models. Relative seed set was calculated by dividing individual seed set by the mean seed set of the replicate.

Our fitness variable had a zero-inflated distribution because ca. half of the plants in each replicate did not produce any seeds. Therefore, we chose to analyze selection through two separate models: a binary response model and a linear “truncated regression” model. The binary response model assessed selection across treatment and among all experimental plants, whereas the truncated model estimated selection only among plants that produced seeds. Fitness in the first model is a binary variable, with 0 for plants that did not produce any seeds and 1 for plants that produced seeds. In the truncated model, we used relative seed set as fitness component, which was truncated to be > 0. Selection gradients were compared between treatments in a pairwise way and computed from the contrasts between treatments with emmeans post-hoc tests (R package *emmeans*: (Lenth 2021)). All statistical analyses were performed with R software 4.0.0 (2020, R Foundation for Statistical Computing, Vienna, Austria).

## Results

### Phenotypic plasticity of traits

Our *B. rapa* plants showed strong phenotypic plasticity, with both morphology and floral volatile emission being affected by our herbivory- and soil treatments (Figure 1, 2), however, no significant interactions between the effects of herbivory and soil were detected (Table 2). Reaction norms (Figure S2 and S3) showed that the full-sib seed family’s responses were relatively homogenous for morphological traits, especially for mPC3 and mPC4 whereas the responses were more heterogenous for floral volatiles (i.e. scent).

**Table 2:**
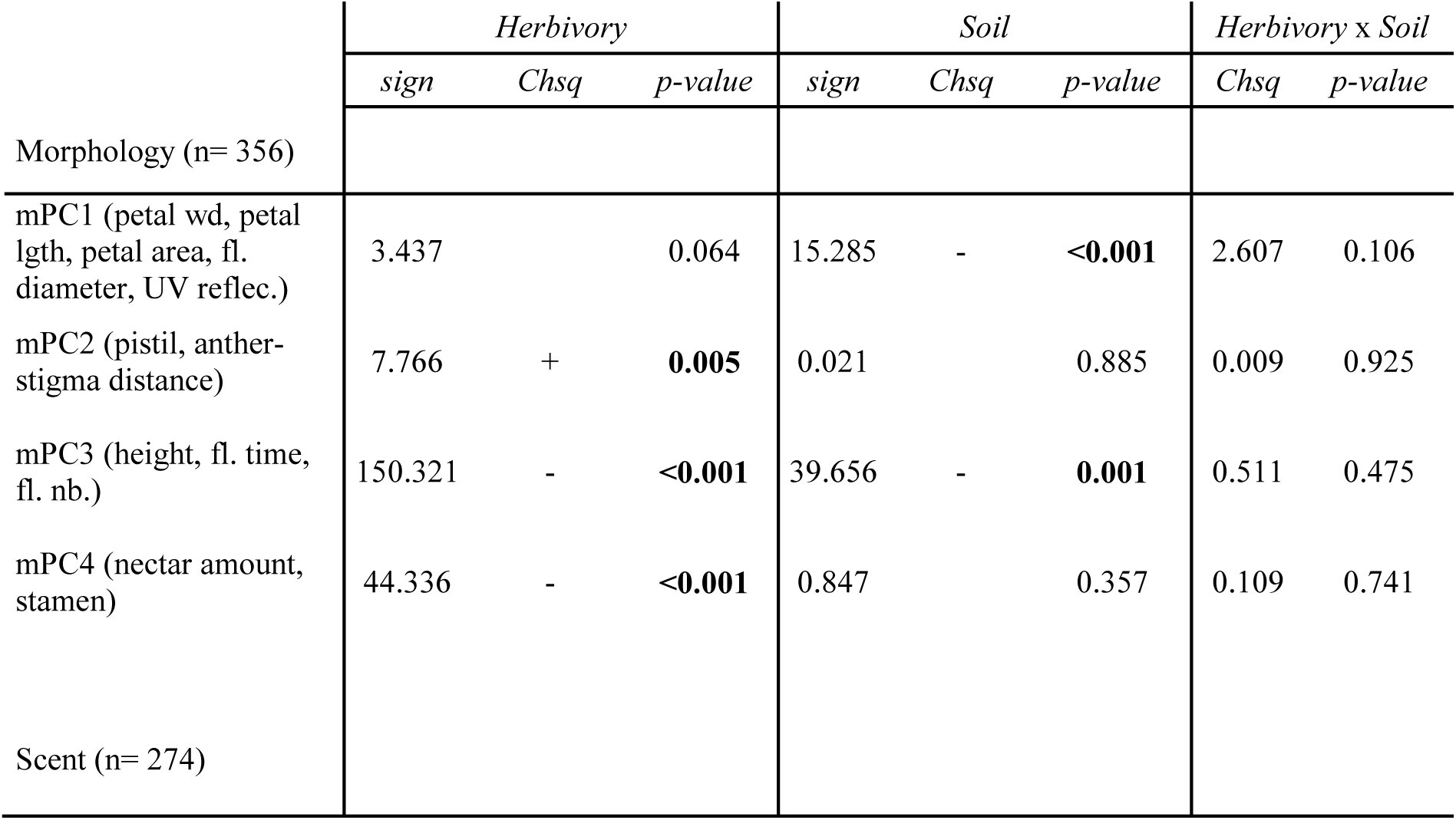
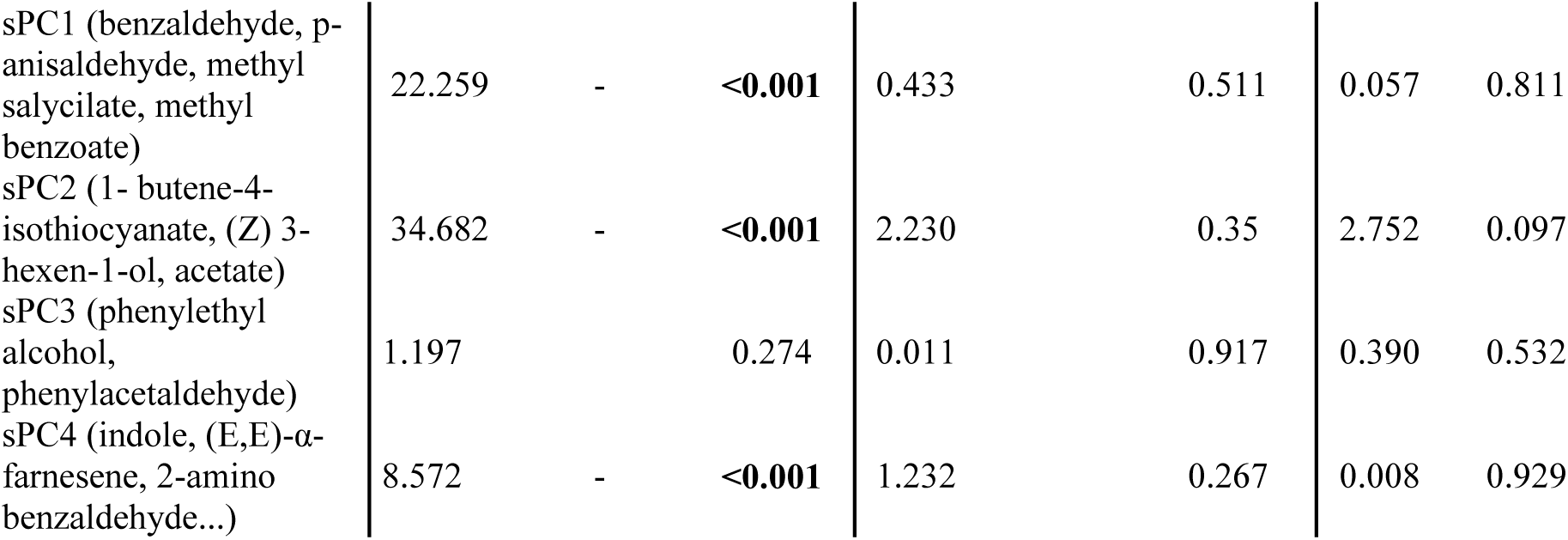
Plant responses to herbivory and soil treatments (i.e. phenotypic plasticity). Principle components (PCs) are shown with the three variables having the highest loadings (see Table 1) indicated in brackets (for responses of individual traits see Table S4 and S5). Signs indicate the difference of means between “no herbivory” versus “herbivory”, and “tuff” versus “limestone” soil. Significant p-value (p<0.05) are given in bold.

**Figure 1:**
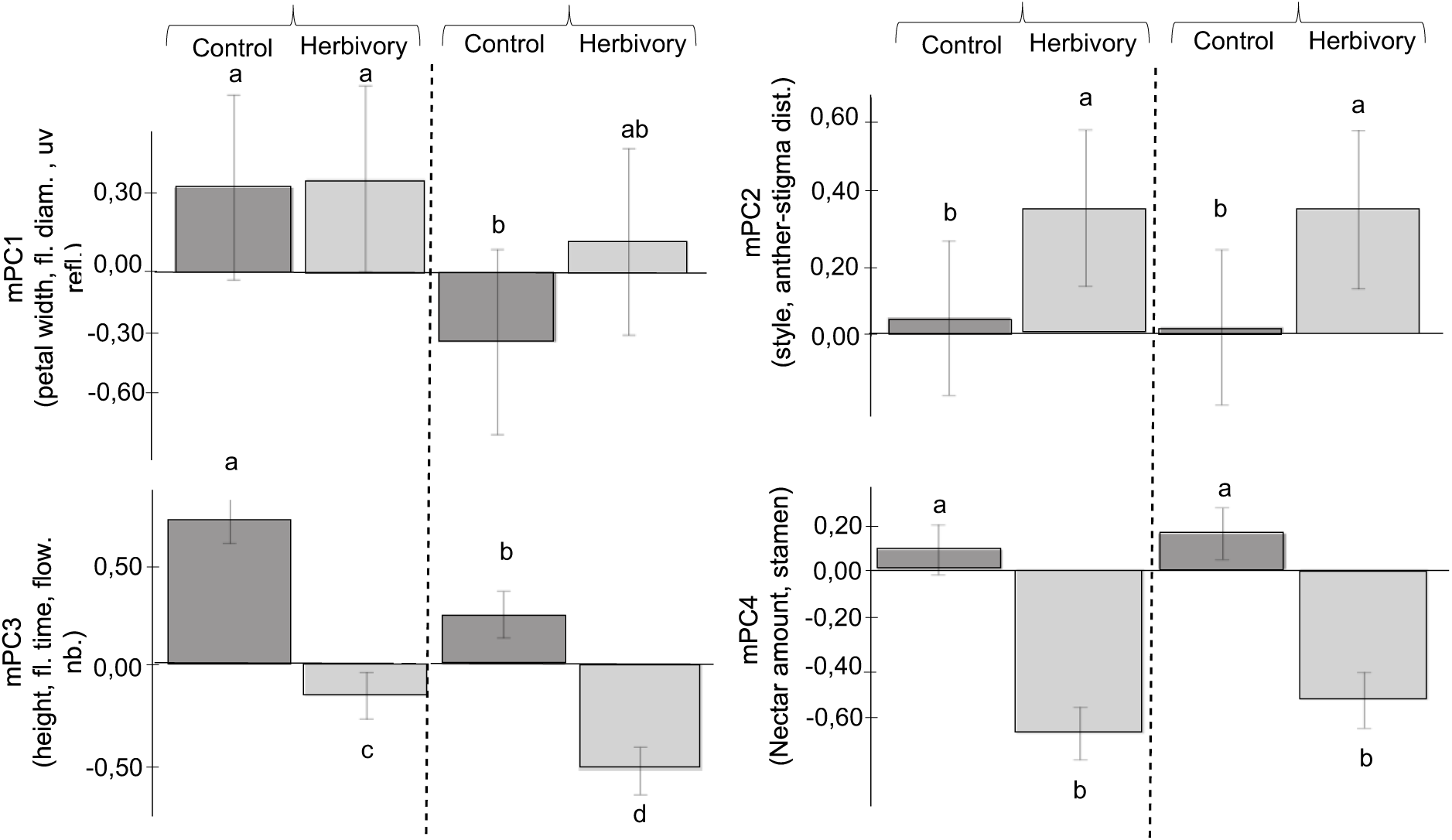
Phenotypic plasticity in *Brassica rapa* morphology with different soil and herbivory. The figure shows how mean ± s.e.m. values of morphological traits (shown as principle components, mPCs) change in response to soil and herbivory. mPC1 (petal width, petal length, petal area, flower diameter, UV-reflecting area), of open flowers), mPC2 (anther-stigma distance, pistil length), mPC3 (plant height, flowering time, number and mPC4 (nectar amount (μL) and stamen length), Significant differences (p<0.05) between treatments are denoted by different letters.

**Figure 2:**
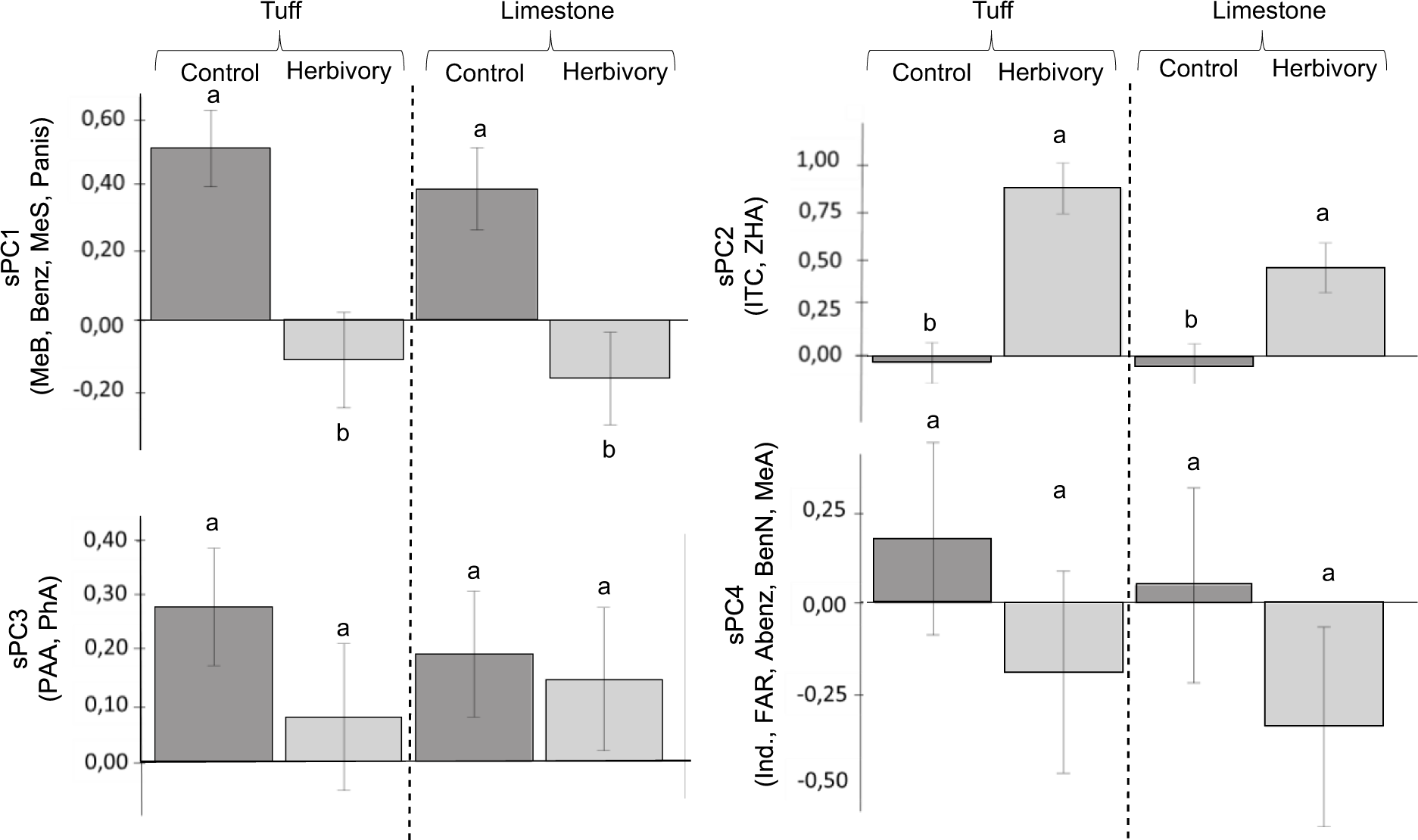
Phenotypic plasticity in *Brassica rapa* floral scent with different soil and herbivory. The figure shows how mean ± s.e.m. values of floral volatiles (i.e. scent, shown as principle components, sPCs) change in response to soil and herbivory. sPC1 (MeB: methyl benzoate, MeS: methyl salicylate,, Benz: benzaldehyde, Panis: p-anisaldehyde), sPC2 (ITC: 1-butene-4-isothiocyanate, ZHA: (Z) 3-hexen-1-ol, acetate), sPC3 (PAA: phenylacetaldehyde, PhA: phenylethyl alcohol) and sPC4 (Ind: indole, FAR: (E,E)-α-farnesene, MeA, methyl anthranilate, Abenz: amino benzaldehyde, BenN: Benzyl nitrile). Significant differences (p<0.05) between treatments are denoted by different letters.

Aphid herbivory induced strong responses in most of the studied traits, with only scent PCs 3 and 4 not being significantly affected (Table 2 and Figures 1 and 2). Plant responses were particularly strong for morphology PC 3 and 4 (Table 2 and Figure 1), indicating that herbivory reduced plant height and flower numbers, delayed flowering (mPC3), lowered nectar production as well as reduced stamen (mPC4) and increased anther-stigma distance (mPC2). In terms of floral volatiles, herbivory reduced the emission of sPC1 (benzaldehyde, p-anisaldehyde, methyl benzoate and methyl salicylate), and increased sPC2 (1-butene-4-isothiocyanate, (*Z*)-3-hexen-1-ol, acetate; Table 2 and Figure 2).

Soil type, on the other hand, only had significant effects on morphology but did not affect floral scent. Generally, plants did much better in tuff soil, with earlier flowering, taller plants, and higher number of flowers (mPC3). As a herbivore-specific effect, we found larger flowers (mPC1) in tuff compared to limestone soil only in non-herbivory plants (Figure 1).

### Resource limitation

We detected resource limitation in plants growing in both soil types, as the addition of fertilizer increased seed set in plants in both soils (Figure S1). Hand-pollinated plants growing in tuff soil produced higher number of seeds, seeds per silique, weight of seeds and plant biomass than plants growing in limestone soil (Figure S1, Table S3), suggesting resource limitation was stronger in limestone soil than in tuff soil.

### Phenotypic selection

We found significant selection on morphological traits and nectar, but not on floral scent. For the binary model including all plants (hand- and bee-pollinated, 1337 plants) significant selection was detected for mPC3 and mPC4, indicating selection for taller plants, greater number of open flowers, earlier flowering (mPC3), longer stamen, and higher amount of nectar (mPC4; Table 3). We also found significant interactions of mPC3 x pollination, mPC3 x pollination x soil type, mPC3 x soil type x herbivory, and mPC3 x pollination x soil type x herbivory (Table 3). The truncated regression model detected also mPC3 and mPC4 under significant selection, and a significant interaction between mPC3 x soil type x herbivory; results for truncated models are reported in Table S7 (all plants), Table S8 (plants with hand pollination) and Table S9 (plants with bee pollination). Because of the clear effect of pollination on selection, and to ease the interpretation of the results, we split the data into plants with hand- and bee-pollination respectively, and analyzed selection separately.

**Table 3:**
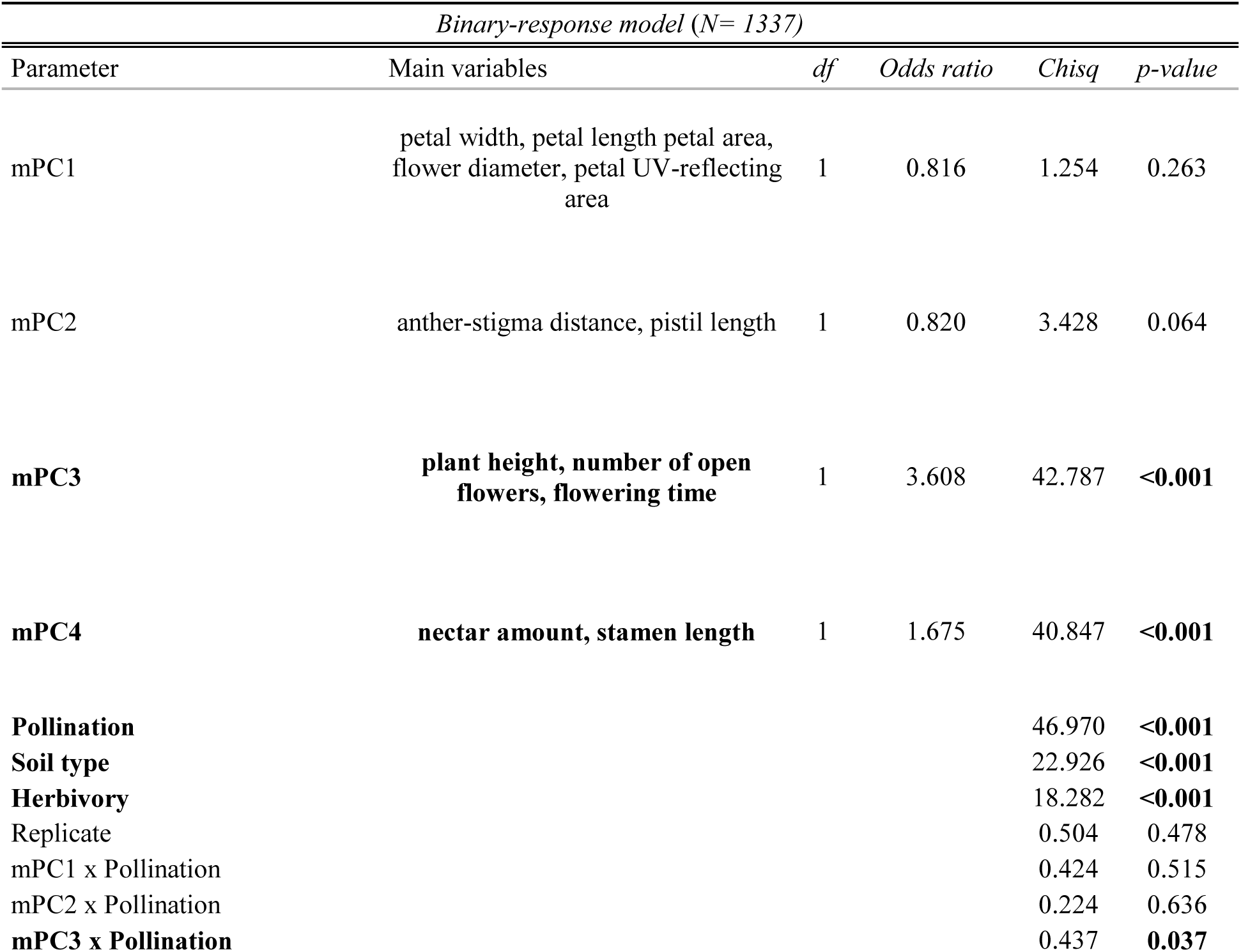
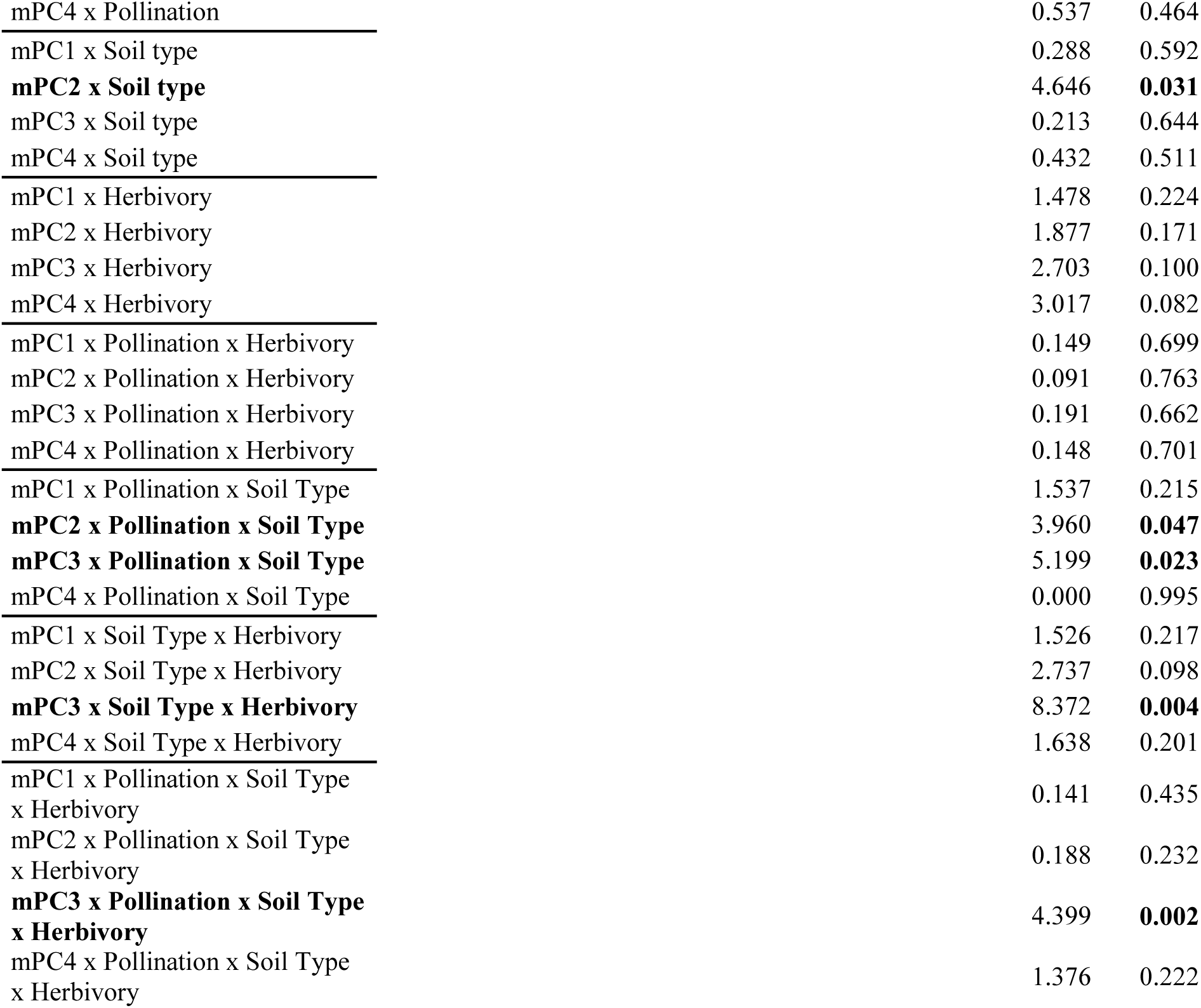
Phenotypic selection in **all plants** (hand- and bee-pollinated) on plant morphology (morphology principal components, mPCs) as estimated by binary generalized linear model with fitness (seeds/no seeds) as dependent variable. Plant morphology principle components (mPCs) were used covariates, pollination, herbivory and soil type as fixed factors, and replicate as random factor. For all plants included (N=1337), fitness is expressed as a binary variable, with 731 plants having no seeds (0) and 606 having seeds (seed set ≥1; 1). Multiple-comparison post hocs were ran using estimated marginal means (EMMs) and their contrasts were computed with the emmeans package (Lenth 2021). P-values are Tukey-corrected for multiple comparisons. Significant p-values (p<0.05) are given in bold.

| *Binary-response model (N= 1337)* |  |  |  |  |  |
| --- | --- | --- | --- | --- | --- |
| Parameter | Main variables | df | Odds ratio | Chisq | p-value |
| mPC1 | petal width, petal length petal area, flower diameter, petal UV-reflecting area | 1 | 0.816 | 1.254 | 0.263 |
| mPC2 | anther-stigma distance, pistil length | 1 | 0.820 | 3.428 | 0.064 |
| **mPC3** | **plant height, number of open flowers, flowering time** | 1 | 3.608 | 42.787 | **<0.001** |
| **mPC4** | **nectar amount, stamen length** | 1 | 1.675 | 40.847 | **<0.001** |
| **Pollination** |  |  |  | 46.970 | **<0.001** |
| **Soil type** |  |  |  | 22.926 | **<0.001** |
| **Herbivory** |  |  |  | 18.282 | **<0.001** |
| Replicate |  |  |  | 0.504 | 0.478 |
| mPC1 x Pollination |  |  |  | 0.424 | 0.515 |
| mPC2 x Pollination |  |  |  | 0.224 | 0.636 |
| **mPC3 x Pollination** |  |  |  | 0.437 | **0.037** |

|  |  |  |
| --- | --- | --- |
| mPC4 x Pollination | 0.537 | 0.464 |
| mPC1 x Soil type | 0.288 | 0.592 |
| **mPC2 x Soil type** | 4.646 | **0.031** |
| mPC3 x Soil type | 0.213 | 0.644 |
| mPC4 x Soil type | 0.432 | 0.511 |
| mPC1 x Herbivory | 1.478 | 0.224 |
| mPC2 x Herbivory | 1.877 | 0.171 |
| mPC3 x Herbivory | 2.703 | 0.100 |
| mPC4 x Herbivory | 3.017 | 0.082 |
| mPC1 x Pollination x Herbivory | 0.149 | 0.699 |
| mPC2 x Pollination x Herbivory | 0.091 | 0.763 |
| mPC3 x Pollination x Herbivory | 0.191 | 0.662 |
| mPC4 x Pollination x Herbivory | 0.148 | 0.701 |
| mPC1 x Pollination x Soil Type | 1.537 | 0.215 |
| **mPC2 x Pollination x Soil Type** | 3.960 | **0.047** |
| **mPC3 x Pollination x Soil Type** | 5.199 | **0.023** |
| mPC4 x Pollination x Soil Type | 0.000 | 0.995 |
| mPC1 x Soil Type x Herbivory | 1.526 | 0.217 |
| mPC2 x Soil Type x Herbivory | 2.737 | 0.098 |
| **mPC3 x Soil Type x Herbivory** | 8.372 | **0.004** |
| mPC4 x Soil Type x Herbivory | 1.638 | 0.201 |
| mPC1 x Pollination x Soil Type<br>x Herbivory | 0.141 | 0.435 |
| mPC2 x Pollination x Soil Type<br>x Herbivory | 0.188 | 0.232 |
| **mPC3 x Pollination x Soil Type**<br>**x Herbivory** | 4.399 | **0.002** |
| mPC4 x Pollination x Soil Type<br>x Herbivory | 1.376 | 0.222 |

For plants with hand-pollination (654 plants), we detected selection on mPC3 and mPC4, however, none of the interactions were significant, showing that selection did not differ among the treatments (Table 4, Figure S3, B). For plants with bee-pollination (683 plants), both models detected significant and positive selection on mPC3 and mPC4, as well as a significant interaction between mPC3 and soil and herbivory (Table 5; Table S9), showing that selection for plant height, flowering time, and number of open flowers (mPC3) differed between treatments.

**Table 4:**
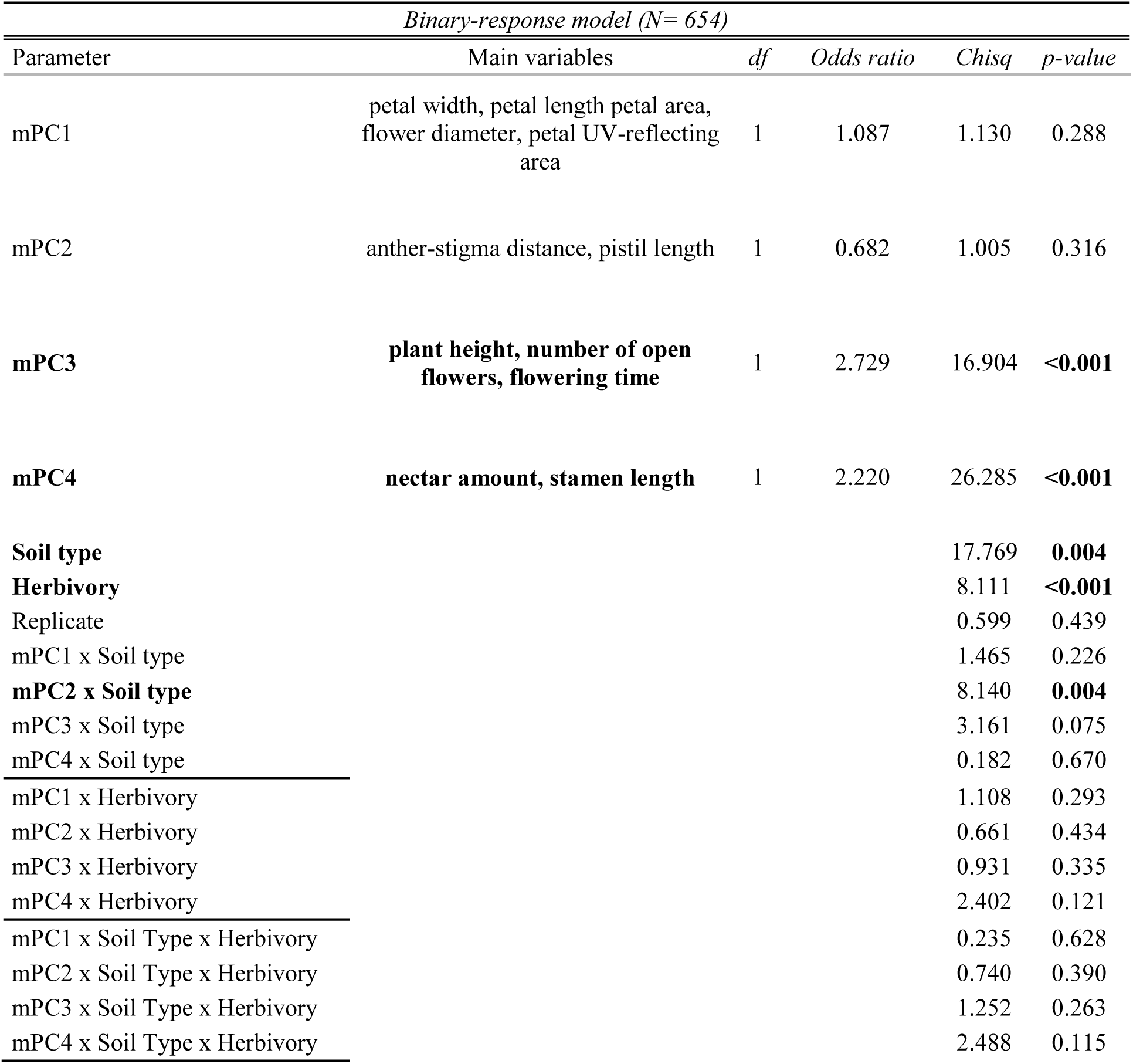
Phenotypic selection in **plants with hand pollination** on plant morphology (morphology principal components, mPCs) as estimated by binary generalized linear model with fitness (seeds/no seeds) as dependent variable. For this analyse, only plants of hand pollination treatments were considered to dissociate phenotypic selection mediated by pollinators to others agents of selection such as herbivore or/ and soil. Plant morphology principle components (mPCs) were used covariates, herbivory and soil type as fixed factors, and replicate as random factor. For all plants included (N=654), fitness is expressed as a binary variable, with 291 plants having no seeds (0) and 363 having seeds (seed set ≥1; 1). Multiple-comparison post hocs were ran using estimated marginal means (EMMs) and their contrasts were computed with the emmeans package (Lenth 2021). P-values are Tukey-corrected for multiple comparisons. Significant p-values (p<0.05) are given in bold.

| *Binary-response model (N= 654)* |  |  |  |  |  |
| --- | --- | --- | --- | --- | --- |
| Parameter | Main variables | df | Odds ratio | Chisq | p-value |
| mPC1 | petal width, petal length petal area, flower diameter, petal UV-reflecting area | 1 | 1.087 | 1.130 | 0.288 |
| mPC2 | anther-stigma distance, pistil length | 1 | 0.682 | 1.005 | 0.316 |
| **mPC3** | **plant height, number of open flowers, flowering time** | 1 | 2.729 | 16.904 | **<0.001** |
| **mPC4** | **nectar amount, stamen length** | 1 | 2.220 | 26.285 | **<0.001** |
| **Soil type** |  |  |  | 17.769 | **0.004** |
| **Herbivory** |  |  |  | 8.111 | **<0.001** |
| Replicate |  |  |  | 0.599 | 0.439 |
| mPC1 x Soil type |  |  |  | 1.465 | 0.226 |
| **mPC2 x Soil type** |  |  |  | 8.140 | **0.004** |
| mPC3 x Soil type |  |  |  | 3.161 | 0.075 |
| mPC4 x Soil type |  |  |  | 0.182 | 0.670 |
| mPC1 x Herbivory |  |  |  | 1.108 | 0.293 |
| mPC2 x Herbivory |  |  |  | 0.661 | 0.434 |
| mPC3 x Herbivory |  |  |  | 0.931 | 0.335 |
| mPC4 x Herbivory |  |  |  | 2.402 | 0.121 |
| mPC1 x Soil Type x Herbivory |  |  |  | 0.235 | 0.628 |
| mPC2 x Soil Type x Herbivory |  |  |  | 0.740 | 0.390 |
| mPC3 x Soil Type x Herbivory |  |  |  | 1.252 | 0.263 |
| mPC4 x Soil Type x Herbivory |  |  |  | 2.488 | 0.115 |

**Table 5:** Phenotypic selection in **plants with bumblebee-pollination** on plant morphology (morphology principal components, mPCs) as estimated by binary generalized linear model with fitness (seeds/no seeds) as dependent variable. Plant morphology principle components (mPCs) were used as covariates, treatment as fixed factor, and replicate as random factor. For all plants included (N=683), fitness is expressed as a binary variable, with 430 plants having no seeds (0) and 253 having seeds (seed set ≥1; 1). Multiple-comparison post hocs were ran using estimated marginal means (EMMs) and their contrasts were computed with the emmeans package (Lenth 2021). P-values are Tukey-corrected for multiple comparisons. Significant p-values (p<0.05) are given in bold.

| Binary-response model (N= 683) |  |  |  |  |  |
| --- | --- | --- | --- | --- | --- |
| Parameter | Main variables | df | Odds ratio | Chisq | p-value |
| mPC1 | petal width, petal length petal area, flower diameter, petal UV-reflecting area | 1 | 0.812 | 0.216 | 0.642 |
| mPC2 | anther-stigma distance, pistil length | 1 | 0.821 | 2.650 | 0.104 |
| mPC3 | plant height, number of open flowers, flowering time | 1 | 3.690 | 28.069 | <0.001 |
| mPC4 | nectar amount, stamen length | 1 | 1.677 | 14.376 | <0.001 |
| Soil type |  |  |  | 10.597 | <0.001 |
| Herbivory |  |  |  | 6.560 | 0.010 |
| Replicate |  |  |  | 0.058 | 0.809 |
| mPC1 x Soil type |  |  |  | 0.187 | 0.666 |
| mPC2 x Soil type |  |  |  | 0.037 | 0.848 |
| mPC3 x Soil type |  |  |  | 1.513 | 0.219 |
| mPC4 x Soil type |  |  |  | 0.171 | 0.679 |
| mPC1 x Herbivory |  |  |  | 0.504 | 0.478 |
| mPC2 x Herbivory |  |  |  | 1.314 | 0.252 |
| mPC3 x Herbivory |  |  |  | 2.182 | 0.140 |
| mPC4 x Herbivory |  |  |  | 0.783 | 0.376 |
| mPC1 x Soil Type x Herbivory |  |  |  | 1.428 | 0.232 |
| mPC2 x Soil Type x Herbivory |  |  |  | 2.319 | 0.128 |
| mPC3 x Soil Type x Herbivory |  |  |  | 9.760 | 0.002 |
| mPC4 x Soil Type x Herbivory |  |  |  | 0.008 | 0.929 |
| Post Hoc tests |  |  |  |  |  |
| Parameter | Treatment groups | n | z - value | p-value |  |
| mPC3 (plant height, number of open flowers, flowering time) | LH - LNH | 312 | 3.814 | 0.001 |  |
|  | LH - TH | 360 | 4.369 | <0.001 |  |
|  | LH - TNH | 349 | 4.719 | <0.001 |  |
|  | LNH - TNH | 323 | 0.621 | <0.001 |  |
|  | LNH - TH | 334 | 4.410 | 0.925 |  |
|  | TH - TNH | 371 | 3.769 | 0.001 |  |

We found no significant selection on any of the floral scent principle components (Tables S10, S11). This may have been a consequence of the lower sample size in the scent data set. Some plants were simply too small to fit into the scent collection glass cylinders and thus their scent could not be sampled.

### Differences in selection in bumblebee-pollinated plants

The significant interaction between mPC3 soil type and herbivory showed that bumblebee-mediated selection in plants differed depending on soil and herbivore attack. We found significant and positive selection on mPC3 in all treatments except in plants in limestone soil without herbivory (LNH; Table 6). In terms of differences in selection among treatment groups, selection on plants growing in limestone soil with herbivory (LH) was significantly stronger than selection in all other treatments (Table 5, Figure 3). On the contrary, plants in tuff showed a greater selection on mPC3 when herbivores were absent (Table 5, Figure 3).

**Table 6:** Phenotypic selection in plants with bumblebee-pollination on plant morphology PC3 estimated individually for the different treatment groups (N=679) with replicate as random factor, by binary generalized linear models with fitness as dependent variable. Fitness is expressed as a binary variable: 0 no seeds (N=426) 1: seed set (N=253). Significant p-values (p<0.05) are given in bold.

| *Morphology (binary-response model)* |  |  |  |  |  |  |  |
| --- | --- | --- | --- | --- | --- | --- | --- |
| Trait | Treatment |  | n | df | Odds Ratio | Chsq | p-value |
| PC3: Height, Flowering Time, Flower number | TNH | Tuff soil, no herbivory | 191 | 1 | 3.059 | 4.779 | **<0.001** |
|  | TH | Tuff soil, herbivory | 180 | 1 | 3.052 | 4.979 | **0.026** |
|  | LNH | Limestone soil, no herbivory | 169 | 1 | 1.154 | 0.663 | 0.416 |
|  | LH | Limestone soil, herbivory | 143 | 1 | 3.052 | 15.114 | **<0.001** |

**Figure 3:**
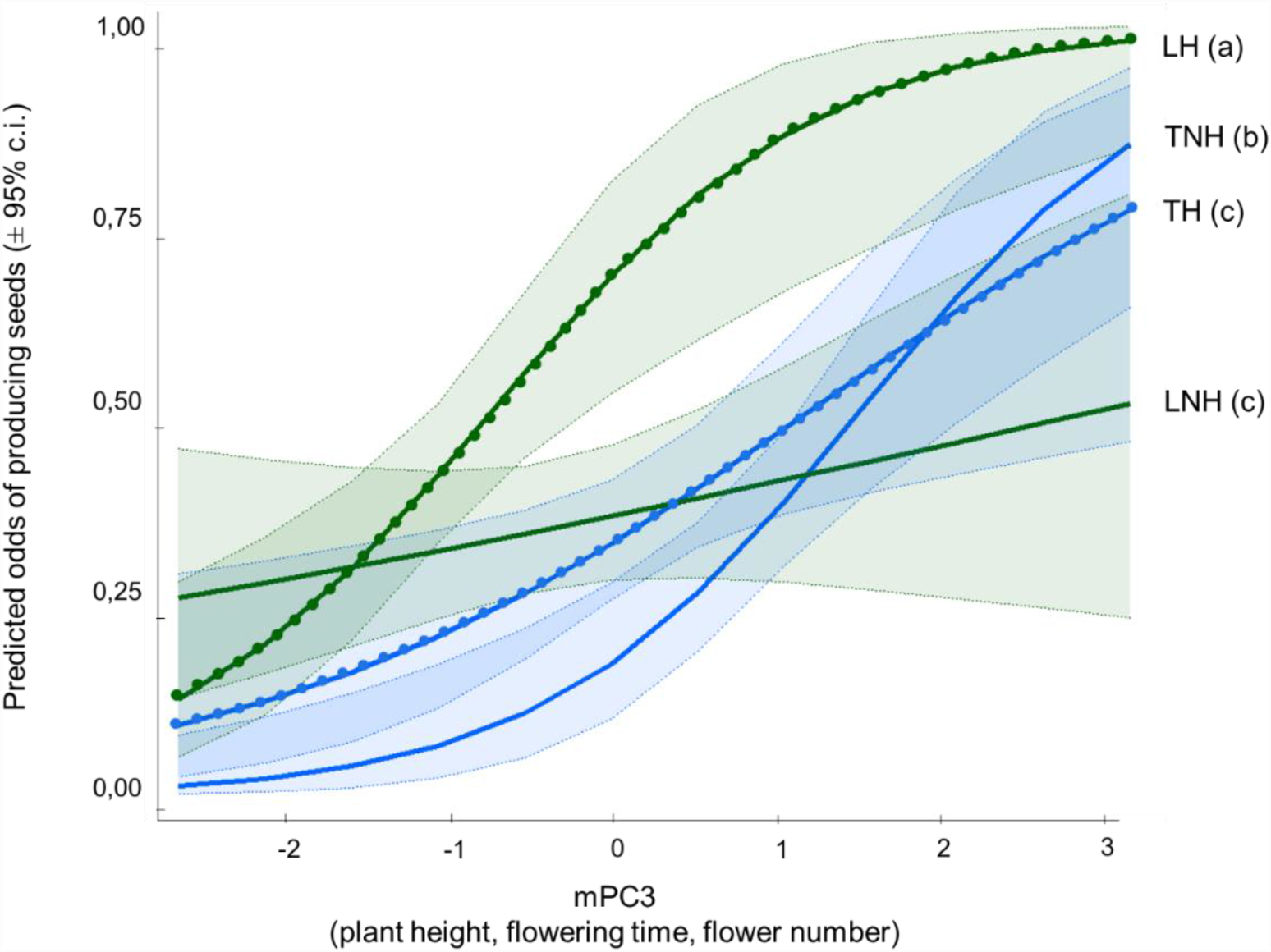
Selection on mPC3 (plant height, flowering time, number of open flowers) in bee-pollinated plants of the different treatment groups: TNH (tuff no herbivory, blue solid line), TH (tuff herbivory, blue dotted line), LNH (limestone no herbivory, green solid lines) and LH (limestone herbivory, green dotted line). Shaded areas denote the 95% confidence interval. Different letters indicate slopes that are significantly different across treatments (p<0.05), as calculated using estimated marginal means (EMMs) and their contrasts using the R package emmeans.

Patterns of bumblebee visitation and seed set differed for plants in limestone soil with (LH) or without herbivory (LNH; Figure 4), despite overall seed production not being different (Table S3). To analyze and visualize this, mPC3 was categorized into 5 categories with 1 representing smallest plants with fewest flowers, and 5 tallest plants with most flowers (see method section). In the treatment with herbivory (LH), tall plants (category 5) had a higher visitation rate and seed production (Figure 4; visits: F_1.59_=4.92, p-value =0.03; seeds: F_1.59_=6.8, p-value =0.012). Total visits were significantly and positively associated with mPC3 categories for LH (β=0.224, p-value=0.007) but not for LNH (β=0.055; p-value =0.475); total seed set was significantly associated with mPC3 in both treatments, but stronger so for LH (β=0.303, p-value <0.001) than for LNH (β=0.158, p-value =0.041).

**Figure 4.**
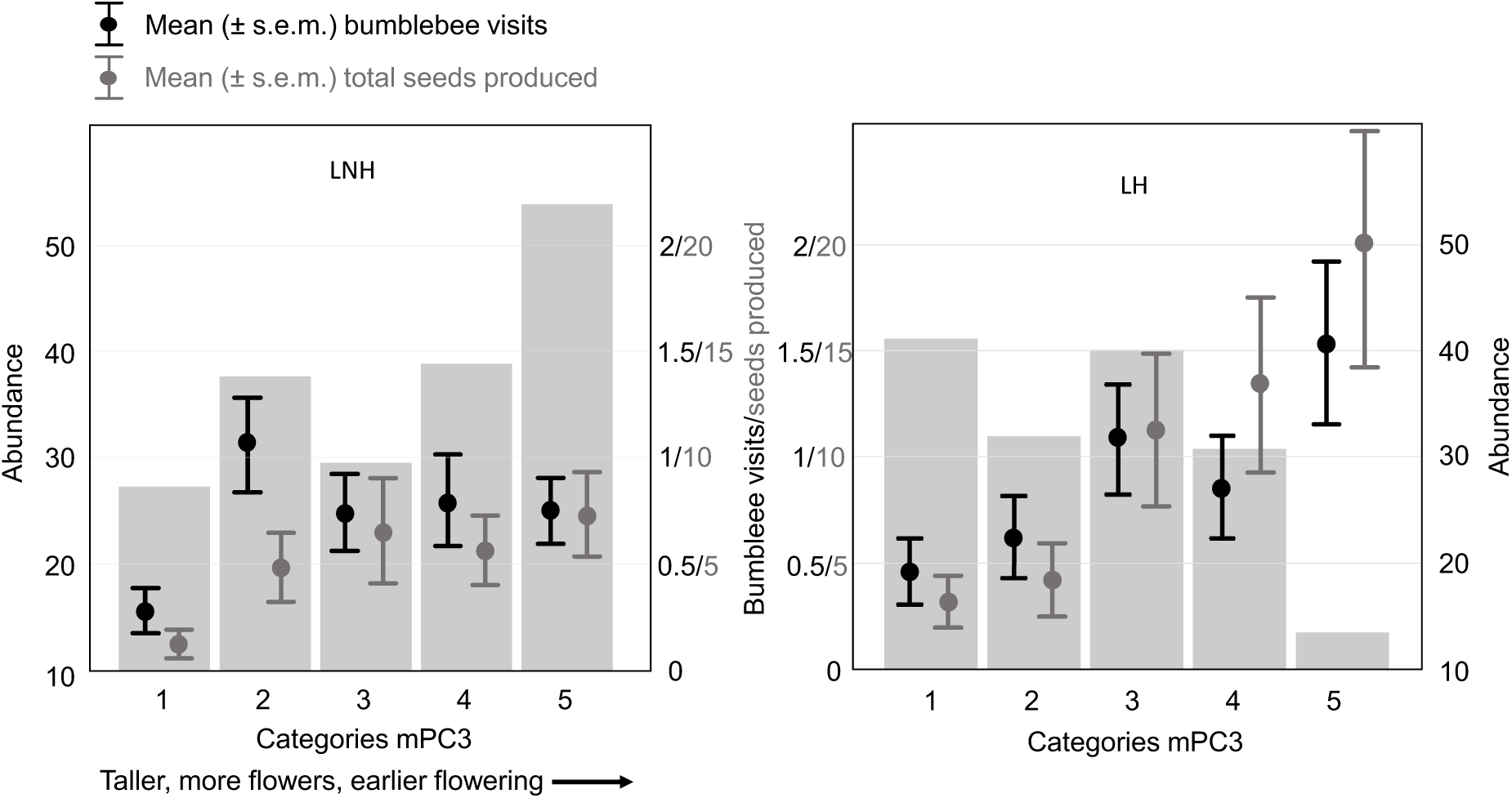
Abundance of plants of different size, flower number and flowering time (mPC3) and their attractiveness to bumblebees in two treatments with limestone soil. The histograms show the distribution of plants falling in five categories of mPC3 for the treatments limestone non-herbivory (LNH, left) and limestone herbivory (LH, right). The black error bars show the mean visitation by bumblebees, and the grey error bars the mean total seeds set of the plants in the five categories. In LH, the (few) tall plants with many flowers (cat. 5) get more visits and produce more seeds than plants of the same category in LNH (see results section for stats) leading to an over-proportionally high seed production (grey error bars), and thus stronger selection for high values of mPC3. Sample size, LNH: category 1: 25, 2:34, 3:27, 4:35, 5:48; total: 169; LH: 1:37, 2: 29, 3: 36, 4:28, 5:13; total: 143.

## Discussion

Phenotypic plasticity is thought to influence adaptive evolution in several different ways, for example through release of otherwise cryptic genetic variation, genetic accommodation, or through changing patterns of selection (West-Eberhard 1989, Pigliucci et al. 2006, Levis and Pfennig 2021). In the latter context, adaptive plasticity is thought to reduce selection and hence slow down adaptive evolution, whereas maladaptive plasticity would have the opposite effect (Hendry 2016, Pfennig 2021). An interesting twist to this idea is the possibility of plasticity to alter the way organisms interact with each other, and therefore to change (reciprocal) selection mediated by the interaction (Agrawal 2001, Fordyce 2006, Hendry 2016). Given that interactions are ubiquitous in nature, and the manifold ways plasticity may alter them, this connection between plasticity and evolution may be common in nature.

In principle, environmental parameters such as soil and herbivores can influence natural selection through two mechanisms. First, these factors may impact the resources available for seed production, through nutrient availability in soil or removal of plant tissue by herbivory (McCall and Irwin 2006, Smallegange et al. 2007, Shi et al. 2013, Lucas-Barbosa 2016). Under resource limitation, pollination-mediated selection acting on seed production (female fitness) is thought to be weaker, because not all fertilized ovules develop into seeds, and thus even the most successfully pollinated plants may not produce more seeds than those with fewer pollination success (Totland 2001, Maad and Alexandersson 2004, Caruso et al. 2005). Second, by modifying pollinator-attracting signals and rewards, these two factors can change the behavior of pollinators and thus modify selection (Strauss et al. 2002, Burkle and Irwin 2009, Zangerl and Berenbaum 2009). Naturally, such indirect effects may not be limited to plant-pollinator interactions, but concern the whole ecological interaction network an organism is engaged in.

Very few studies, however, have quantified the consequences of plasticity-modified interactions on phenotypic selection and/or adaptive evolution. Recently, Ramos and Schiestl (2019) have shown, using experimental evolution, that leaf herbivory can impact pollinator-driven evolution. This study also documented different patterns of selection with and without herbivory, e.g. for herkogamy, likely caused by herbivore-induced phenotypic changes in plants.

Our study detects selection mediated by resource limitation, as shown in the plants with hand pollination. In these plants, the same principle components were under selection as in plants with bee-pollination. However, those PCs showed no significant interactions with either soil type or herbivory, suggesting the environmental factors did not impact selection differently across treatments. Indeed resource limitation was detected for both soil types, however, with a tendency to increase in limestone soil. A significant interaction between PC3 and the experimental factors was only apparent for bumblebee-pollinated plants, pointing towards the importance of indirect, interactor-mediated effects causing different patterns of selection.

The way environmental factors could have changed pollinator-mediated selection in our experiment is by “producing” different phenotypes, thus altering the signals that bees use when choosing plants for visitation. Both soil and aphids could have hardly impacted the bee’s choices directly, as soil surface was very similar for both soil types and aphids were removed before pollination. Both soil type and herbivory, however, caused strong changes in the plant’s phenotypes. For soil, the likely reason is the different chemical environment and nutrient availability to the plants. Tuff- and limestone soil differ largely in their CaCO_3_ concentration (tuff soil: 6.0g/100g of soil; limestone soil: 19.4g/100g). Calcareous soils such as those formed by limestone are characterized by an excess of carbonate ions (CaCo3>15%; Alloway 2008) along with a high pH. In these alkaline conditions, most of the essential nutrients are under a form that is unavailable to plants, which leads to nutrient deficiencies. The most common and well documented limitations are N, P, Zn, Fe deficiencies (Marschner 1995, Alloway 2008), which have negative effects on growth rates and may cause “stress phenotypes” (Agarwala et al. 1988, Shi et al. 2013).

Herbivory is well known to induce changes in plant morphology and volatile emission. We found that herbivory reduced primarily the production of aromatic volatiles, such as methyl benzoate and p-anisaldehyde, two bumblebee attractants (Schiestl et al. 2014, Gervasi and Schiestl 2017). Similar results were previously shown for *Brassica rapa* with chewing herbivores (Schiestl et al. 2014) and other *Brassicaceae* with aphids (Pareja et al. 2012, Rusman et al. 2019). Aromatic compounds are biosynthetically derived from phenylalanine through the enzyme phenylalanine ammonia-lyase (PAL), which is also implicated in plant defense against aphids (Eck et al. 2010), suggesting a trade-off between volatile production and aphid-induced defense. Other floral traits that were negatively affected by aphid feeding and may face similar trade-offs or be affected by resource-reallocation were flowering time, plant height, nectar amount, all known to affect the attractiveness of plants to pollinators (Agrawal 1999a, Cipollini 2010, Schiestl et al. 2014).

Aphid-induced plasticity affected phenotypic selection on plants in both soil types, but differently so. While for plants in (richer) tuff soil, selection on plant size/flower number was stronger without herbivory, but the opposite was found for plants in (poorer) limestone soil. This interactive effect was particularly obvious for limestone soil, where we detected a change in bumblebee behavior in response to infested and non-infested plants. In limestone soil, plants with aphid herbivory faced the most stressful environment, and commonly remained small (shown in the left histogram of Figure 4). Bees seemingly discriminated more strongly between tall and short plants in this treatment (Figure 4), leading to an over-proportional visitation and seed production by tall plants, causing stronger selection on height and flower number (i.e. high values of mPC3).

Our study shows that the combination of two environmental factors modify phenotypic selection in an interactive way. In our experiment, the behavior of the main selective agent, the pollinator, was the most important causative agents of this change. We argue that variable selection caused by phenotypic plasticity may be common in nature, but its effect on adaptive evolution will depend on the consistency of plastic change and the ecological factors causing plasticity and selection, as well as population genetic parameters such as evolvability. This study, focusing on plasticity and selection, has been followed up by an evolution experiment, using the same experimental parameters through 10 generations, to document the evolutionary changes in plants of the eight treatment groups, the results of which will be published in subsequent papers. We feel that the link between phenotypic plasticity, selection, and adaptive evolution is particularly fascinating and deserves more attention in future research.

## Acknowledgements

We sincerely thank Luca Arrigo, Giovanni Scopece, Léa Frachon for their help with soil collection and phenotyping. We further thank Laura Dällenbach for her help with measurements of plant traits and insect rearing, Rayko Jonas and Markus Meierhofer for their help with growing plants, Franz Huber for his help with gas chromatographic analyses, and Michael Whitlock for fruitful discussions on the manuscript. The research was funded by the Swiss National Science Funds (SNF grant no. 31003A_172988 to FPS). Data will be archived upon acceptance at the University of Zürich Zora open repository archive.

