## Supplemental data for "Plant phenotypic plasticity changes pollinator-mediated selection"

### Supplementary Information

**Table S1** – Physical, chemical, and mineralogical properties of each soil used in this study. Soil physics and chemistry was analyzed by the INRA laboratories following the SOL-1031 protocol (Table S1)(Ciesielski et al. 1997). This method used a solution of HF-HLCO4, an acid mixed which allowed to digest soil clay and thus quantify both available and unavailable nutrients to plants.

| | | | Limestone | Tuff |
|---|---|---|---|---|
| Physical parameters | Clay | % | 9.30 | 6.70 |
| | Fine Silt | | 17.40 | 13.70 |
| | Coarse Silt | | 6.90 | 12.80 |
| | Fine Sand | | 14.20 | 20.80 |
| | Coarse Sand | | 52.20 | 46.00 |
| Chemical Parameters | N | g/kg | 2.57 | 1.06 |
| | P | g/kg | 3.61 | 1.79 |
| | Org. Carbon | g/kg | 28.40 | 12.50 |
| | Soil organic matter | g/kg | 49.10 | 21.60 |
| | C/N | | 11.00 | 11.80 |
| | Cu | g / 100 g | 0.48 | 0.35 |
| | Zn | g / 100 g | 0.77 | 0.91 |
| | CaCO3 | g / 100 g | 19.40 | 0.60 |
| | Si | g / 100 g | 17.00 | 26.50 |
| | Ca | meq / 100 g | 5.94 | 1.12 |
| | Mg | meq / 100 g | 0.91 | 0.26 |
| | K | meq / 100 g | 0.75 | 1.56 |
| | Na | meq / 100 g | 0.37 | 0.96 |
| | Fe | meq / 100 g | 5.59 | 4.05 |
| | Mn | meq / 100 g | 9.93 | 11.71 |
| | Al | meq / 100 g | 0.31 | 8.53 |

**Table S2** – Rotated loadings on the principal components of morphology and scent combined for *Brassica rapa* in generation one and two. Only principal components with eigenvalues> 0.9 and that explain> 4 % of the total variance were retained.

| | PC1 | PC2 | PC3 | PC4 | PC5 | PC6 |
|---|---|---|---|---|---|---|
| Eigen Values | 6.56 | 4.14 | 1.84 | 1.48 | 1.29 | 1.10 |
| Variance explained (%) | 26.23 | 16.57 | 7.35 | 5.91 | 5.17 | 4.39 |
| Rotated loading of components | | | | | | |
| Plants height | 0.23 | 0.27 | **0.77** | -0.05 | -0.06 | 0.17 |
| Flowering time | -0.14 | -0.16 | **-0.75** | 0.02 | -0.16 | 0.15 |
| Flower number | -0.06 | 0.08 | **0.77** | -0.03 | -0.19 | 0.18 |
| Nectar amount (μL) | 0.13 | 0.17 | 0.05 | -0.11 | 0.07 | **0.70** |
| Pistil | 0.06 | 0.27 | -0.03 | 0.07 | **0.81** | 0.25 |
| Stamen | 0.14 | 0.32 | -0.06 | -0.04 | -0.07 | **0.70** |
| Anther- stigma distance | -0.01 | 0.03 | -0.06 | 0.07 | **0.88** | -0.13 |
| Petal width | 0.08 | **0.76** | 0.20 | -0.06 | 0.04 | 0.22 |
| Petal length | 0.11 | **0.67** | 0.13 | -0.06 | 0.26 | 0.50 |
| Petal area | 0.06 | **0.92** | 0.13 | 0.07 | 0.04 | -0.10 |
| Uv reflecting area | -0.02 | **0.87** | 0.06 | 0.16 | 0.01 | -0.03 |
| Flower diameter | 0.05 | **0.67** | 0.12 | -0.13 | 0.23 | 0.44 |
| Benzaldehyde | 0.44 | -0.06 | -0.28 | **0.54** | -0.21 | 0.05 |
| 1- butene-4- isothiocyanate | -0.02 | 0.04 | 0.09 | **0.70** | 0.16 | -0.14 |
| Methyl benzoate | **0.55** | -0.07 | 0.17 | 0.43 | 0.00 | 0.30 |
| Phenylethyl alcohol | **0.72** | 0.24 | 0.07 | 0.17 | -0.07 | -0.12 |
| 2-Amino benzaldehyde | **0.87** | 0.01 | 0.04 | -0.10 | 0.03 | 0.10 |
| p-Anisaldehyde | **0.58** | -0.02 | -0.07 | 0.32 | -0.12 | 0.14 |
| Methyl anthranilate | **0.80** | -0.10 | 0.09 | 0.09 | 0.02 | 0.11 |
| (Z) 3-Hexen-1-ol. acetate | 0.21 | 0.09 | -0.11 | **0.62** | 0.04 | -0.14 |
| Phenylacetaldehyde | **0.75** | -0.11 | 0.02 | -0.01 | -0.04 | -0.03 |
| Benzyl nitrile | **0.87** | 0.07 | 0.15 | 0.00 | 0.04 | 0.09 |
| Methyl salicylate | **0.59** | -0.10 | 0.15 | 0.42 | -0.08 | 0.26 |
| Indole | **0.76** | 0.04 | -0.08 | 0.04 | 0.10 | 0.04 |
| (E. E ) -α- Farnesene | **0.53** | 0.00 | 0.13 | 0.31 | 0.31 | 0.09 |

Variables in bold with loading>0.50 were considered significant

**Table S3** Fitness values (mean ± s.d.) of plants of the differnet treatment groups (T=tuff, L=limestone, NH=no herbivory, H=herbivory) with bumblebee and hand pollination. Statistical differences (ANOVA with LSD posthoc test, $P<0.05$) are indicated by different letters. Sample size is 196 plants per treatment.

| | *Bee pollination (N=98)* | | | | *Hand pollination (N=98)* | | | |
|---|---|---|---|---|---|---|---|---|
| | TNH | TH | LNH | LH | THN | TH | LNH | LH |
| Total number of visits | $0.72^{A}$ ± 1.04 | $0.71^{A}$ ± 1.04 | $0.72^{A}$ ± 1.07 | $0.71^{A}$ ± 1.19 | | | | |
| Time per visit (s) | $61.52^{A}$ ± 15.48 | $35.28^{B}$ ± 94.49 | $44.18^{AB}$ ± 82.69 | $29.53^{B}$ ± 74.99 | | | | |
| Time per flower (s) | $8.29^{A}$ ± 17.87 | $5.02^{B}$ ± 11.67 | $8.96^{AC}$ ± 15.44 | $3.51^{B}$ ± 9.00 | | | | |
| Number of silliques | $1.84^{A}$ ± 3.04 | $1.6^{AC}$ ± 2.46 | $1.03^{B}$ ± 1.73 | $1.18^{BC}$ ± 2.09 | $1.74^{A}$ ± 1.81 | $1.88^{A}$ ± 1.83 | $1.57^{A}$ ± 1.71 | $1.66^{A}$ ± 1.73 |
| Number of seeds | $12.21^{A}$ ± 23.27 | $10.84^{AC}$ ± 21.25 | $5.08^{B}$ ± 10.51 | $7.49^{BC}$ ± 15.76 | $20.96^{A}$ ± 24.64 | $22.19^{A}$ ± 26.59 | $14.90^{B}$ ± 18.44 | $13.22^{B}$ ± 17.04 |
| Seeds per sillique | $6.32^{A}$ ± 3.04 | $6.26^{A}$ ± 4.04 | $4.62^{B}$ ± 2.90 | $6.76^{A}$ ± 3.98 | $11.68^{A}$ ± 5.13 | $11.67^{A}$ ± 8.02 | $9.10^{B}$ ± 3.82 | $7.33^{C}$ ± 3.93 |
| Seed weight (g) | $0.02^{A}$ ± 0.04 | $0.02^{A}$ ± 0.03 | $0.01^{B}$ ± 0.02 | $0.01^{B}$ ± 0.02 | $0.03^{A}$ ± 0.04 | $0.03^{A}$ ± 0.04 | $0.02^{B}$ ± 0.02 | $0.02^{B}$ ± 0.02 |

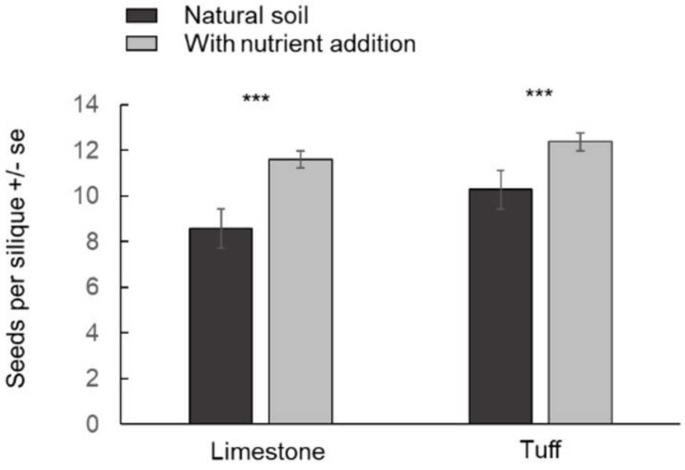


**Figure S1:** Variation in mean seed number per silique in plants with hand pollination growing in the two soil types before and after addition of fertilizer. Significance was determined using a generalized mixed model with seeds per silique as dependent variable, "fertilizer" and "soil type" as fixed factors, their interaction, and replicate as random factors. The data shows that the use of fertilizer led to a significant increase in seed production (effect of "fertilizer", $Chi^2$: 20.561, p<0.001), and thus seed production is resource-limited in both soils. Model also detected a tendency for tuff to produce more seeds (effect of "soil", $Chi^2$: 3.322, p=0.068) suggesting that seeds production in limestone may be more resource-limited than in tuff. There was no significant interaction detected in the model (effect of "fertilizer x Soil", $Chi^2$: 0.964, p=0.405).

**Table S4** – Plant responses to herbivory and soil treatments in generation 1. Signs are indicated by comparing no herbivory versus herbivory and limestone compared to tuff soil. Significant p-value (p<0.05) are given in bold.

| | *Herbivory* | *Soil* | *Herbivory* x *Soil* |
|---|---|---|---|

| | *sign* | *Chsq* | *p-value* | *sign* | *Chsq* | *p-value* | *Chsq* | *p-value* |
|---|---|---|---|---|---|---|---|---|
| Morpho | *0-1* | | | *L-T* | | | | |
| mPC1 | | 3.437 | 0.064 | - | 15.285 | **<0.001** | 2.607 | 0.106 |
| mPC2 | + | 7.766 | **0.005** | | 0.021 | 0.885 | 0.009 | 0.925 |
| mPC3 | - | 150.321 | **<0.001** | - | 39.656 | **0.001** | 0.511 | 0.475 |
| mPC4 | - | 44.336 | **<0.001** | | 0.847 | 0.357 | 0.109 | 0.741 |
| Plants height | - | 266.397 | **<0.001** | + | 15.271 | **<0.001** | 3.652 | 0.056 |
| Flowering time | + | 70.107 | **<0.001** | - | 63.479 | **<0.001** | 1.333 | 0.248 |
| Flower number | - | 22.270 | **<0.001** | + | 8.252 | **0.004** | 3.233 | 0.072 |
| Nectar amount (μL) | - | 35.719 | **<0.001** | | 1.368 | 0.242 | 0.533 | 0.465 |
| Pistil | | 1.624 | 0.203 | | 0.006 | 0.981 | 0.522 | 0.470 |
| Stamen | - | 45.852 | **<0.001** | | 0.254 | 0.614 | 0.002 | 0.961 |
| Anther- stigma distance | + | 17.548 | **<0.001** | | 0.037 | 0.848 | 0.260 | 0.610 |
| Petal width | | 0.173 | 0.677 | + | 56.663 | **<0.001** | 2.036 | 0.154 |
| Petal length | - | 9.898 | **0.001** | + | 9.261 | **0.002** | 0.779 | 0.378 |
| Petal area | | 0.771 | 0.380 | + | 10.133 | **0.001** | 4.331 | **0.037** |
| Uv reflecting area | | 0.309 | 0.578 | | 2.833 | 0.092 | 2.386 | 0.122 |
| Flower diameter | - | 8.588 | **0.003** | + | 18.719 | **<0.001** | 1.947 | 0.163 |
| Scent | | | | | | | | |
| sPC1 | - | 22.259 | **<0.001** | | 0.433 | 0.511 | 0.057 | 0.811 |
| sPC2 | - | 34.682 | **<0.001** | | 2.230 | 0.350 | 2.752 | 0.097 |
| sPC3 | | 1.197 | 0.274 | | 0.011 | 0.917 | 0.390 | 0.532 |
| sPC4 | - | 8.572 | **<0.001** | | 1.232 | 0.267 | 0.008 | 0.929 |
| Benzaldehyde | | 2.590 | 0.108 | | 0.196 | 0.658 | 0.040 | 0.842 |
| 1- butene-4-isothiocyanate | + | 28.571 | **<0.001** | + | 5.705 | **0.017** | 2.294 | 0.130 |
| Methyl benzoate | - | 28.304 | **<0.001** | | 3.342 | 0.068 | 0.465 | 0.452 |
| Phenylethyl alcohol | - | 4.060 | **0.044** | | 2.172 | 0.141 | 0.559 | 0.455 |
| 2-Amino benzaldehyde | - | 25.044 | **<0.001** | | 1.186 | 0.172 | 0.249 | 0.618 |
| p-Anisaldehyde | - | 6.044 | **0.014** | | 0.321 | 0.571 | 0.364 | 0.547 |
| Methyl anthranilate | - | 19.947 | **<0.001** | | 0.722 | 0.396 | 0.355 | 0.551 |
| (Z) 3-Hexen-1-ol, acetate | | 2.346 | 0.126 | | 0.890 | 0.345 | 0.590 | 0.443 |
| Phenylacetaldehyde | - | 8.938 | **0.003** | | 0.077 | 0.782 | 0.001 | 0.979 |
| Benzyl nitrile | - | 24.710 | **<0.001** | | 3.437 | 0.064 | 0.004 | 0.984 |
| Methyl salicylate | - | 15.281 | **<0.001** | | 1.777 | 0.183 | 0.036 | 0.849 |
| Indole | - | 8.188 | **0.004** | | 0.036 | 0.850 | 0.283 | 0.595 |
| (E,E)-α-Farnesene | | 2.369 | 0.124 | | 0.311 | 0.577 | 0.070 | 0.791 |
| Total VOCs | - | 12.615 | **<0.001** | | 1.295 | 0.255 | 0.535 | 0.464 |

**Table S5** – Trait differences (mean ± s.d.) between plants across treatment groups at generation 1. Treatments differ by the soil in which plants grow (L: limestone; T: tuff) and by herbivory

(CO: control, H: with herbivory). All scent compounds are expressed in pg flower $l^{-1}$ sampled air$^{-1}$, units of other traits are indicated in the table.

| Morphology | TNH | TH | LNH | LH |
|---|---|---|---|---|
| mPC1 | 0.31 ± 1.05$^{A}$ | 0.33 ± 0.98$^{A}$ | -0.24 ± 0.92$^{B}$ | 0.09 ± 1.01$^{AB}$ |
| mPC2 | 0.04 ± 1.01$^{A}$ | 0.34 ± 1.03$^{B}$ | 0.01 ± 0.93$^{A}$ | 0.34 ± 1.25$^{B}$ |
| mPC3 | 0.73 ± 0.63$^{A}$ | -0.16 ± 0.74$^{C}$ | 0.25 ± 0.77$^{B}$ | -0.54± 0.93$^{D}$ |
| mPC4 | 0.59 ± 0.82$^{A}$ | -0.59 ± 1.05$^{B}$ | 0.13 ± 0.58$^{A}$ | -0.47 ± 0.65$^{B}$ |
| Height (cm) | 28.40 ± 7.12$^{A}$ | 17.82 ± 6.29$^{C}$ | 25.06 ± 4.64$^{B}$ | 16.63 ± 5.26$^{C}$ |
| Flowering Time (day) | 21.50 ± 1.66$^{C}$ | 23.42 ± 2.65$^{B}$ | 23.32 ± 2.58$^{B}$ | 25.85 ± 3.05$^{A}$ |
| Flower number | 6.85 ± 4.12$^{A}$ | 4.63 ± 3.14$^{B}$ | 5.29 ± 3.15$^{B}$ | 4.27 ± 2.60$^{B}$ |
| Nectar (μL/flower) | 76.29 ± 44.74$^{A}$ | 47.73 ± 34.52$^{B}$ | 74.78 ± 37.35$^{A}$ | 52.52 ± 39.72$^{B}$ |
| Pistil (cm) | 0.86 ± 0.13 | 0.69 ± 0.14 | 0.67 ± 0.13 | 0.70 ± 0.16 |
| Stamen (cm) | 0.63 ± 0.07$^{A}$ | 0.56 ± 0.09$^{B}$ | 0.61 ± 0.06$^{A}$ | 0.56 ± 0.09$^{B}$ |
| Anther- stigma distance (cm) | 0.132 ± 0.09$^{B}$ | 0.17 ± 0.09$^{A}$ | 0.13 ± 0.08$^{B}$ | 0.18 ± 0.11$^{A}$ |
| Petal width (cm) | 0.40 ± 0.05$^{A}$ | 0.39 ± 0.06$^{A}$ | 0.35 ± 0.05$^{B}$ | 0.36 ± 0.86$^{B}$ |
| Petal length (cm) | 0.92 ± 0.09$^{A}$ | 0.88 ± 0.09$^{B}$ | 0.88 ± 0.09$^{B}$ | 0.86 ± 0.09$^{B}$ |
| Petal area ($cm^2$) | 0.25 ± 0.08$^{A}$ | 0.23 ± 0.06$^{AB}$ | 0.21 ± 0.06$^{B}$ | 0.22 ± 0.07$^{B}$ |
| Uv reflecting area ($cm^2$) | 0.20 ± 0.07 | 0.19 ± 0.06 | 0.18 ± 0.06 | 0.19 ± 0.07 |
| Flower diameter (cm) | 13.10 ± 1.31$^{A}$ | 12.53 ± 1.31$^{B}$ | 12.35 ± 1.25$^{B}$ | 12.14 ± 1.20$^{B}$ |
| **Scent** | | | | |
| sPC1 | 0.50 ± 1.14$^{A}$ | -0.12 ± 1.04$^{B}$ | 0.38 ± 1.02$^{A}$ | -0.18 ± 0.79$^{B}$ |
| sPC2 | -0.04 ± 1.00$^{B}$ | 0.87 ± 1.04$^{A}$ | -0.05 ± 0.89$^{B}$ | 0.46 ± 1.00$^{A}$ |
| sPC3 | 0.27 ± 1.10 | 0.08 ± 0.86 | 0.19 ± 1.00 | 0.14 ± 0.65 |
| sPC4 | 0.18 ± 1.00 | -0.21 ± 1.02 | 0.09 ± 1.00 | -0.33 ± 1.07 |
| Benzaldehyde | 762.55 ± 664.60 | 592.42 ± 329.63 | 709.86 ± 513.42 | 587.16 ± 369.78 |
| 1-butene 4-isothiocyanate | 69.11 ± 82.12$^{B}$ | 209.00 ± 269.63$^{A}$ | 68.78 ± 513.42$^{B}$ | 102.94 ± 119.11$^{AB}$ |
| Methyl benzoate | 66.06 ± 72.71$^{A}$ | 32.39 ± 56.25$^{B}$ | 51.89 ± 80.68$^{A}$ | 20.47 ± 21.17$^{B}$ |
| Phenylethyl alcohol | 15.61 ± 25.44 | 8.30 ± 13.24 | 11.67 ± 22.59 | 5.73 ± 3.35 |
| 2-Amino benzaldehyde | 749.40 ± 118.32$^{A}$ | 318.99 ± 318.99$^{BC}$ | 572.51 ± 821.71$^{AB}$ | 145.91 ± 233.86$^{C}$ |
| p-Anisaldehyde | 42.30 ± 57.60 | 25.39 ± 29.85 | 33.05 ± 30.49 | 20.52 ± 24.40 |
| Methyl anthranilate | 52.90 ± 73.21$^{A}$ | 15.47 ± 23.18$^{B}$ | 50.17 ± 81.07$^{AB}$ | 16.84 ± 41.25$^{B}$ |
| 3-Hexen-1-ol, acetate (Z) | 43.470 ± 69.16 | 75.14 ± 170.28 | 41.43 ± 70.06 | 36.82 ± 55.76 |
| Phenylacetaldehyde | 265.13 ± 494.40 | 109.61 ± 25.82 | 215.10 ± 454.25 | 64.77 ± 90.41 |
| Benzyl nitrile | 57.15 ± 88.06$^{A}$ | 20.02 ± 27.40$^{BC}$ | 40.00 ± 56.43$^{AB}$ | 12.32 ± 12.73$^{C}$ |
| Methyl salicylate | 33.31 ± 47.72$^{A}$ | 19.74 ± 23.60$^{AB}$ | 24.96 ± 24.93$^{A}$ | 12.62 ± 13.06$^{B}$ |
| Indole | 232.74 ± 384.25 | 11.66 ± 184.10 | 184.56 ± 305.24 | 98.03 ± 128.04 |

| | | | | |
|---|---|---|---|---|
| (E,E)-α-Farnesene | 280.47 ± 278.56 | 225.09 ± 271.15 | 227.79 ± 191.06 | 203.36 ± 221.46 |
| Total volatiles | 2670.18 ± 2380.00$^{A}$ | 1632.86 ± 970.69$^{A}$ | 2231.76 ± 1662.04$^{AB}$ | 1327.48 ± 632.22$^{BC}$ |

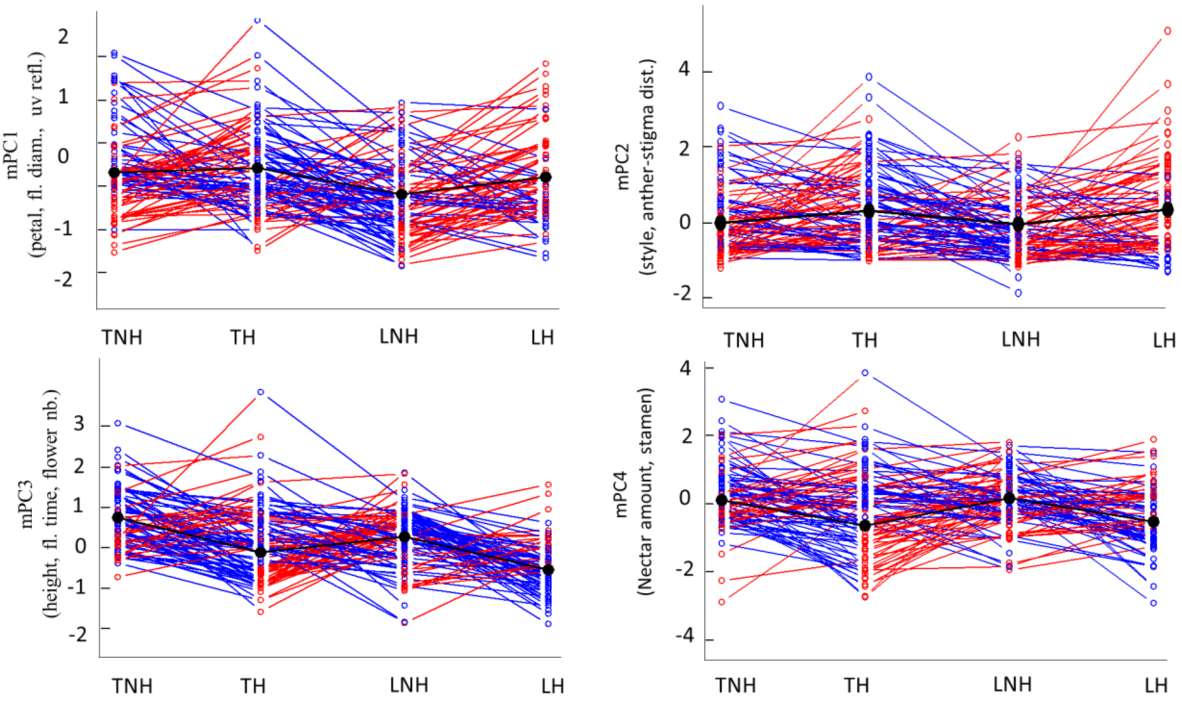


**Figure S2:** Reaction norms showing mPCs scores across treatments. Treatments differ by the soil in which plants grow (L: limestone; T: tuff) and by herbivory (NH: no herbivory, H: with herbivory). Each line represents a sib-seed family (N= 98). Red lines describe positive reaction norms whereas blue describes a negative reaction norm.

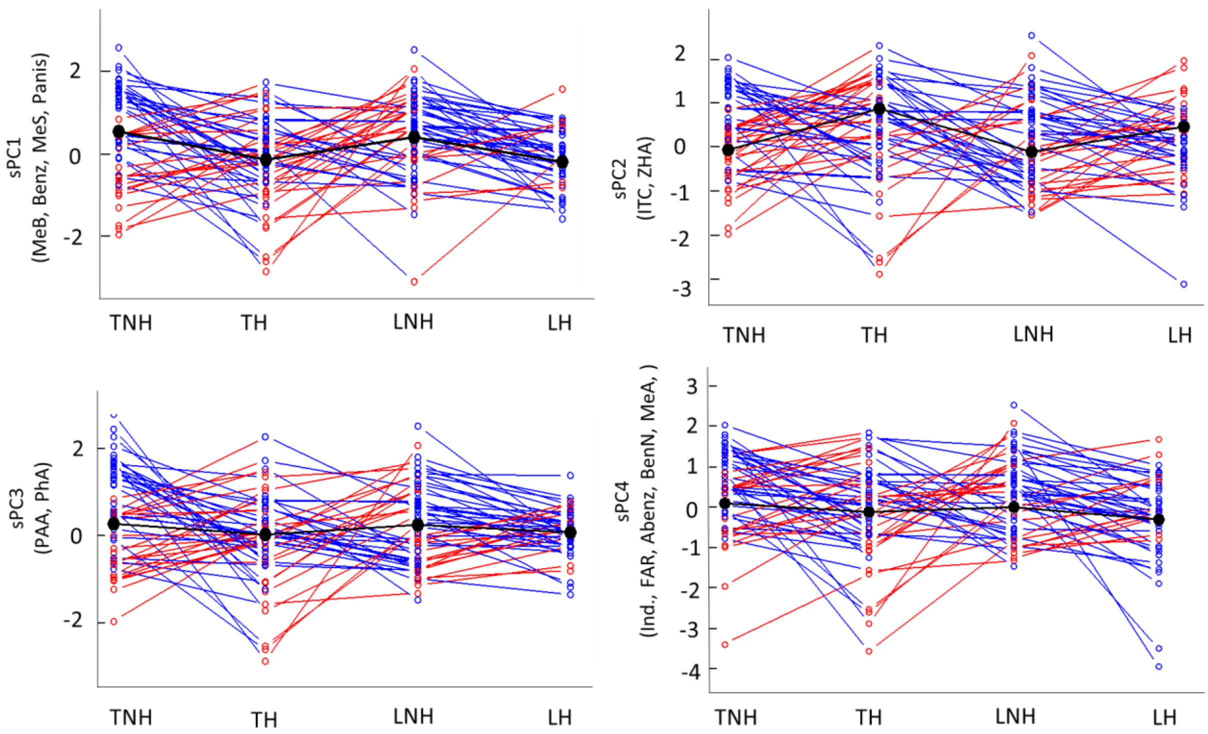


**Figure S3:** Reaction norms showing sPCs scores across treatments. Treatments differ by the soil in which plants grow (L: limestone; T: tuff) and by herbivory (NH: no herbivory, H: with herbivory). Each line represents a sib-seed family (N= 78). Red lines describe positive reaction norms whereas blue describes a negative reaction norm.

**Table S6:** Pearson correlation coefficient of each trait with nectar per flower, mPC3 (principal component represented mainly by nectar and stamen) and visitation time, calculated for plants of all treatments, except for visitation time where only visited plants were considered. (values in bold: $p<0.05$; n.a.: not analysed).

| | Nectar | mPC4 | Visitation Time |
|---|---|---|---|
| Traits | Corr | Corr | Corr |
| Plants height | **0.18** | **0.16** | **0.20** |
| Flowering time | -0.02 | **0.27** | **-0.07** |
| Flower number | **0.11** | **0.21** | **0.32** |
| Nectar amount (µL) | 1.00 | **0.62** | 0.05 |
| Pistil | **0.22** | **0.28** | -0.04 |

| | | | |
|---|---|---|---|
| Stamen | **0.35** | **0.75** | **0.14** |
| Anther- stigma distance | -0.02 | **-0.15** | -0.04 |
| Petal width | **0.24** | 0.04 | **0.15** |
| Petal length | **0.37** | **0.38** | **0.13** |
| Petal area | **0.26** | 0.04 | **0.10** |
| UV reflecting area | **0.14** | 0.00 | 0.02 |
| Flower diameter | **0.37** | **0.32** | **0.09** |
| mPC1 | **0.21** | -0.03 | 0.06 |
| mPC2 | 0.06 | -0.01 | **-0.08** |
| mPC3 | 0.04 | -0.07 | **0.20** |
| mPC4 | **0.62** | 1.00 | **0.14** |
| Benzaldehyde | -0.03 | 0.04 | **0.09** |
| 1- Butene-4- isothiocyanate | **-0.08** | **-0.11** | 0.00 |
| Methyl benzoate | **0.19** | **0.14** | 0.02 |
| Phenylethyl alcohol | **0.12** | **0.11** | **0.10** |
| 2-Amino benzaldehyde | **0.21** | **0.17** | 0.06 |
| p-Anisaldehyde | **0.08** | **0.10** | 0.04 |
| Methyl anthranilate | **0.12** | **0.09** | 0.02 |
| (Z) 3-Hexen-1-ol, acetate | -0.06 | -0.06 | **0.09** |
| Phenylacetaldehyde | **0.21** | **0.17** | 0.07 |
| Benzyl nitrile | **0.21** | **0.16** | 0.03 |
| Methyl salicylate | **0.14** | **0.14** | 0.03 |
| Indole | **0.11** | **0.11** | 0.00 |
| (E,E)-α-Farnesene | **0.08** | 0.05 | 0.01 |
| Total VOCs | **0.14** | **0.11** | 0.07 |
| sPC1 | **0.15** | **0.09** | -0.04 |
| sPC2 | **0.07** | **0.10** | 0.05 |
| sPC3 | **0.12** | **0.12** | **0.10** |
| sPC4 | **-0.14** | **-0.14** | 0.05 |
| Relative Seed Set | n.a | n.a | **0.38** |

**Table S7:** Selection in all plants (hand- and bee-pollinated) on plant morphology estimated by a truncated model with fitness (relative seed set) as dependent variable, including only plants that produced seeds. Plant morphology sPCs were used as covariates, pollination, herbivory and soil type as fixed factors, and replicate as random factor. Linear selection gradients (ß + standard error) are shown, significant p-value ($p<0.05$) are given in bold.

| *Truncated regression model (N=606)* | | | | | |
|---|---|---|---|---|---|
| Parameter | Main variables | *df* | *ß ± sem* | *Chsq* | *p-value* |

| | | | | | |
|---|---|---|---|---|---|
| mPC1 | petal width, petal length petal area, flower diameter, petal UV-reflecting area | 1 | 0.376 ± 0.354 | 2.576 | 0.108 |
| mPC2 | anther-stigma distance, pistil length | 1 | 0.092 ± 0.230 | 0.651 | 0.420 |
| **mPC3** | **plant height, number of open flowers, flowering time** | 1 | 0.898 ± 0.340 | 14.068 | **<0.001** |
| **mPC4** | **nectar amount, stamen length** | 1 | 0.529 ± 0.600 | 4.405 | **0.036** |
| **Pollination** | | | | 43.683 | **<0.001** |
| **Soil type** | | | | 3.023 | 0.082 |
| **Herbivory** | | | | 9.846 | **0,002** |
| Replicate | | | | 0.295 | 0,587 |
| mPC1 x Pollination | | | | 0.101 | 0,750 |
| mPC2 x Pollination | | | | 0.863 | 0,353 |
| mPC3 x Pollination | | | | 0.309 | 0,578 |
| mPC4 x Pollination | | | | 2.259 | 0,133 |
| mPC1 x Soil type | | | | 0.013 | 0,397 |
| mPC2 x Soil type | | | | 0.005 | 0,876 |
| mPC3 x Soil type | | | | 0.000 | 0,675 |
| mPC4 x Soil type | | | | 0.173 | 0,248 |
| mPC1 x Herbivory | | | | 0.718 | 0.908 |
| mPC2 x Herbivory | | | | 0.025 | 0.943 |
| mPC3 x Herbivory | | | | 0.176 | 0.998 |
| mPC4 x Herbivory | | | | 1.334 | 0.678 |
| **mPC1 x Pollination x Herbivory** | | | | **3.846** | **0.050** |
| mPC2 x Pollination x Herbivory | | | | 0.148 | 0.701 |
| mPC3 x Pollination x Herbivory | | | | 0.076 | 0.783 |
| mPC4 x Pollination x Herbivory | | | | 0.306 | 0,580 |
| mPC1 x Pollination x Soil Type | | | | 0.334 | 0.563 |
| mPC2 x Pollination x Soil Type | | | | 0.146 | 0.702 |
| mPC3 x Pollination x Soil Type | | | | 1.045 | 0.307 |
| mPC4 x Pollination x Soil Type | | | | 1.724 | 0.189 |
| mPC1 x Soil Type x Herbivory | | | | 0.514 | 0.473 |
| mPC2 x Soil Type x Herbivory | | | | 0.729 | 0.393 |
| **mPC3 x Soil Type x Herbivory** | | | | **5.293** | **0.021** |
| mPC4 x Soil Type x Herbivory | | | | 0.084 | 0.772 |
| mPC1 x Pollination x Soil Type x Herbivory | | | | 0.642 | 0.423 |

| | | | | |
|---|---|---|---|---|
| mPC2 x Pollination x Soil Type x Herbivory | | | 0.924 | 0.337 |
| mPC3 x Pollination x Soil Type x Herbivory | | | 0.138 | 0.710 |
| mPC4 x Pollination x Soil Type x Herbivory | | | 0.950 | 0.330 |
| *Post Hoc tests* | | | | |
| Parameter | Treatment groups | *n* | *t - value* | *p-value* |
| mPC3 (plant height, number of open flowers, flowering time) x Soil Type x Herbivory | LH - LNH | 271 | 2.700 | **0.036** |
| | LH - TH | 292 | 3.505 | **0.003** |
| | LH - TNH | 295 | 3.742 | **0.001** |
| | LNH - TNH | 314 | 3.598 | **0.002** |
| | LNH – TH | 311 | -0.789 | 0.859 |
| | TH - TNH | 335 | 2.772 | **0.029** |

**Table S8:** Phenotypic selection in plants with hand-pollination on mPCs estimated by a truncated regression model with fitness (relative seed set) as dependent variable, including only those plants that produced seeds. Plant morphology traits (mPCs) were used as covariates, treatment as fixed factor, and replicate as random factor. Linear selection gradients (*ß + standard error*) are shown, significant p-values (p<0.05) are given in bold.

| *Truncated regression model (N=363)* | | | | | |
|---|---|---|---|---|---|
| Parameter | Main variables | *df* | *ß ± sem* | *Chsq* | *p-value* |
| mPC1 | petal width, petal length petal area, flower diameter, petal UV-reflecting area | 1 | -0.026 ± 0.146 | 1.385 | 0.239 |
| mPC2 | anther-stigma distance, pistil length | 1 | -0.024 ± 0.137 | 0.003 | 0.958 |
| **mPC3** | **plant height, number of open flowers, flowering time** | **1** | **0.228 ± 0.183** | **8.338** | **0.004** |
| mPC4 | nectar amount, stamen length | 1 | -0.014 ± 0.131 | 0.153 | 0.695 |
| Soil type | | | | 0.183 | 0,129 |
| Herbivory | | | | 2.310 | 0,668 |
| Replicate | | | | 0.153 | 0,129 |
| mPC1 x Soil type | | | | 0.222 | 0.695 |
| mPC2 x Soil type | | | | 0.027 | 0.144 |
| mPC3 x Soil type | | | | 2.036 | 0.320 |
| mPC4 x Soil type | | | | 0.014 | 0.672 |
| mPC1 x Herbivory | | | | 2.139 | 0.144 |

| | | |
|---|---|---|
| mPC2 x Herbivory | 0.178 | 0.673 |
| mPC3 x Herbivory | 0.027 | 0.870 |
| mPC4 x Herbivory | 0.988 | 0.320 |
| mPC1 x Soil Type x Herbivory | 0.388 | 0.533 |
| mPC2 x Soil Type x Herbivory | 0.040 | 0.841 |
| mPC3 x Soil Type x Herbivory | 0.728 | 0.393 |
| mPC4 x Soil Type x Herbivory | 0.069 | 0.793 |

**Table S9:** Phenotypic selection in plants with bumblebee-pollination on mPCs estimated by a truncated regression model with fitness (relative seed set) as dependent variable, including only those plants that produced seeds. Plant morphology traits (mPCs) were used as covariates, treatment as fixed factor, and replicate as random factor. Linear selection gradients (*ß* + *standard error*) are shown, significant p-values (p<0.05) are given in bold.

| *Truncated linear regression model (N=243)* | | | | | |
|---|---|---|---|---|---|
| Parameter | Main variables | *df* | *ß ± sem* | *Chsq* | *p-value* |
| mPC1 | petal width, petal length petal area, flower diameter, petal UV-reflecting area | 1 | *0.493* ± *0.383* | 1.485 | 0.223 |
| mPC2 | anther-stigma distance, pistil length | 1 | *0.050* ± *0.319* | 0.083 | 0.363 |
| **mPC3** | **plant height, number of open flowers, flowering time** | 1 | *1.335* ± *0.516* | 7.039 | **0.008** |
| **mPC4** | **nectar amount, stamen length** | 1 | *0.763* ± *0.376* | 5.068 | **0.024** |
| **Soil type** | | | | **7.347** | **0.007** |
| Herbivory | | | | 2.957 | 0.085 |
| Replicate | | | | 0.175 | 0.676 |
| mPC1 x Soil type | | | | 0.538 | 0.463 |
| mPC2 x Soil type | | | | 0.025 | 0.874 |
| mPC3 x Soil type | | | | 0.005 | 0.943 |
| mPC4 x Soil type | | | | 2.186 | 0.139 |
| mPC1 x Herbivory | | | | 1.431 | 0.232 |
| mPC2 x Herbivory | | | | 0.082 | 0.775 |
| mPC3 x Herbivory | | | | 0.013 | 0.908 |
| mPC4 x Herbivory | | | | 0.059 | 0.808 |

| | | |
|---|---|---|
| mPC1 x Soil Type x Herbivory | 0.287 | 0,592 |
| mPC2 x Soil Type x Herbivory | 0.796 | 0,372 |
| **mPC3 x Soil Type x Herbivory** | **4.493** | **0.034** |
| mPC4 x Soil Type x Herbivory | 0.002 | 0,967 |

*Post Hoc tests*

| Parameter | Treatment groups | *n* | *z - value* | *p-value* |
|---|---|---|---|---|
| mPC3 (plant height, number of open flowers, flowering time) | LH-LNH | 110 | 2.579 | 0.051 |
| | **LH - TH** | 114 | 3.158 | **0.010** |
| | **LH - TNH** | 121 | 3.377 | **0.005** |
| | **LNH - TNH** | 129 | 2.953 | **0.018** |
| | LNH - TH | 122 | -0.523 | 0.953 |
| | TH - TNH | 133 | 2.431 | 0.074 |

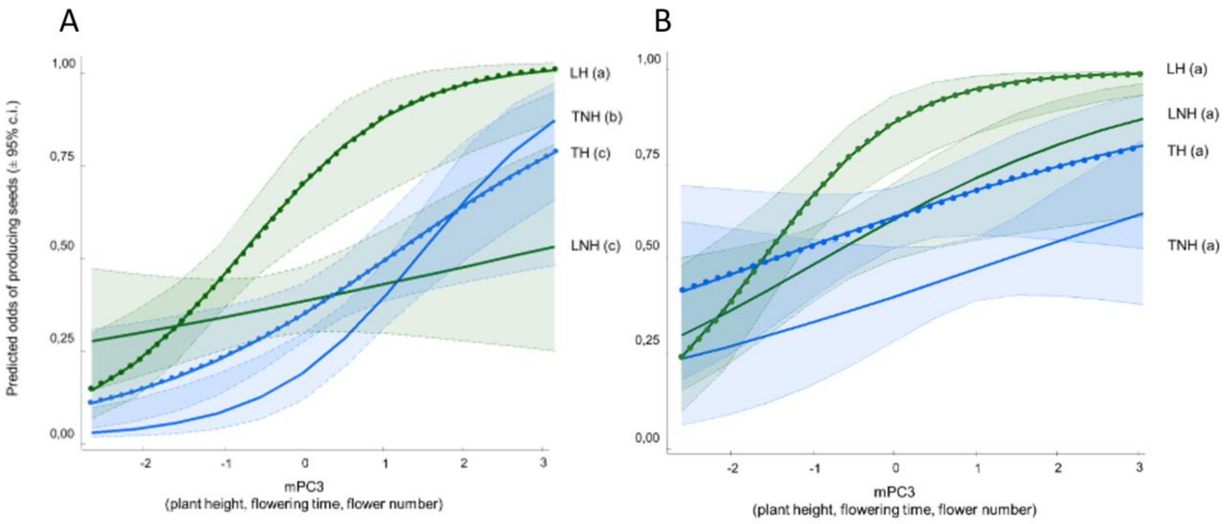


**Figure S3:** Selection on mPC3 (plant height, flowering time, number of open flowers) in plants with bee-pollination (left, A) and hand-pollination (right, B). Treatments differ by the soil in which plants grow (L or T), by herbivory/no herbivory (H or NH). Shaded areas denote the 95% confidence interval. Different letters indicate slopes that are significantly different across treatments ($p<0.05$), as calculated using estimated marginal means (EMMs) and their contrasts using the R package emmeans.

**Table S10:** Phenotypic selection in all plants (hand- and bee-pollinated) on floral volatile emission (scent principal components, sPCs used as covariates) as estimated by binary generalized linear model with fitness (seeds/no seeds) as dependent variable. Floral volatile emission sPCs were used as covariates, pollination, herbivory and soil type as fixed factors, and replicate as random factor. For all plants included (N=1087), fitness is expressed as a binary variable, with 564 plants having no seeds (0) and523 having seeds (seed set ≥1; 1). Significant p-values ($p<0.05$) are given in bold.

| *Binary-response model* (*N= 1087)* | | | | | |
|---|---|---|---|---|---|
| Parameter | Main variables | *df* | *Odds ratio* | *Chisq* | *p-value* |
| sPC1 | methyl benzoate, methyl salycilate, p-anisaldehyde | 1 | 1.403 | 0.103 | 0.749 |
| sPC2 | 1- butene-4- isothiocyanate, (Z) 3-hexen-1-ol, acetate | 1 | 0.790 | 1.787 | 0.181 |
| sPC3 | phenylethyl alcohol, phenylacetaldehyde, 2-amino benzaldehyde, benzyl nitrile | 1 | 0.641 | 1.625 | 0.202 |
| sPC4 | indole, (E,E)-α-farnesene, methyl anthranilate, | 1 | 0.807 | 0.217 | 0.641 |
| **Pollination** | | | | **45.133** | **<0.001** |
| Soil type | | | | 0.029 | 0.865 |
| Herbivory | | | | 0.308 | 0.579 |
| Replicate | | | | 0.007 | 0.931 |
| sPC1 x Pollination | | | | 0.082 | 0.774 |
| sPC2 x Pollination | | | | 1.714 | 0.190 |
| sPC3 x Pollination | | | | 1.441 | 0.230 |
| sPC4 x Pollination | | | | 0.997 | 0.318 |
| sPC1 x Soil type | | | | 0.074 | 0.785 |
| **sPC2 x Soil type** | | | | **5.192** | **0.023** |
| sPC3 x Soil type | | | | 0.149 | 0.699 |
| sPC4 x Soil type | | | | 1.183 | 0.277 |
| **sPC1 x Herbivory** | | | | **14.558** | **<0.001** |
| sPC2 x Herbivory | | | | 0.064 | 0.800 |
| sPC3 x Herbivory | | | | 0.419 | 0.483 |

| | | |
|---|---|---|
| sPC4 x Herbivory | 0.876 | 0.349 |
| sPC1 x Pollination x Herbivory | 0.026 | 0.873 |
| sPC2 x Pollination x Herbivory | 0.216 | 0.612 |
| **sPC3 x Pollination x Herbivory** | **5.443** | **0.020** |
| sPC4 x Pollination x Herbivory | 0.190 | 0.663 |
| sPC1 x Pollination x Soil Type | 0.275 | 0.600 |
| sPC2 x Pollination x Soil Type | 0.042 | 0.838 |
| sPC3 x Pollination x Soil Type | 0.161 | 0.688 |
| sPC4 x Pollination x Soil Type | 0.002 | 0.961 |
| sPC1 x Soil Type x Herbivory | 1.290 | 0.256 |
| sPC2 x Soil Type x Herbivory | 0.761 | 0.383 |
| sPC3 x Soil Type x Herbivory | 0.930 | 0.335 |
| sPC4 x Soil Type x Herbivory | 0.001 | 0.973 |
| sPC1 x Pollination x Soil Type x Herbivory | 1.851 | 0.174 |
| sPC2 x Pollination x Soil Type x Herbivory | 0.015 | 0.904 |
| sPC3 x Pollination x Soil Type x Herbivory | 0.010 | 0.919 |
| sPC4 x Pollination x Soil Type x Herbivory | 1.023 | 0.312 |

**Table S11:** Phenotypic selection in all plants (hand- and bee-pollinated) on floral volatiles estimated by a truncated model with fitness (relative seed set) as dependent variable, including only plants that produced seeds. Floral volatile emission sPCs were used as covariates, pollination, herbivory and soil type as fixed factors, and replicate as random factor. Linear selection gradients (ß + standard error) are shown, significant p-value (p<0.05) are given in bold.

| *Truncated regression model (N=523)* | | | | | |
|---|---|---|---|---|---|
| Parameter | Main variables | *df* | *ß ± sem* | *Chsq* | *p-value* |
| mPC1 | methyl benzoate, methyl salycilate, p-anisaldehyde | 1 | -0.120 ± 0.368 | 0.602 | 0.438 |
| mPC2 | 1- butene-4-isothiocyanate, (Z) 3-hexen-1-ol, acetate | 1 | -0.323 ± 0.295 | 0.057 | 0.812 |
| mPC3 | phenylethyl alcohol, phenylacetaldehyde, 2-amino benzaldehyde, benzyl nitrile | 1 | 1.208 ± 0.376 | 0.177 | 0.674 |

| | | | | | |
|---|---|---|---|---|---|
| mPC4 | indole, (E,E)-α-farnesene, methyl anthranilate, | 1 | -0.166 ± 0.270 | 0.356 | 0.551 |
| **Pollination** | | | | 41.854 | **<0.001** |
| Soil type | | | | 0.030 | 0.862 |
| Herbivory | | | | 0.003 | 0.957 |
| Replicate | | | | 0.399 | 0.528 |
| mPC1 x Pollination | | | | 0.657 | 0.418 |
| mPC2 x Pollination | | | | 1.361 | 0.243 |
| mPC3 x Pollination | | | | 0.026 | 0.872 |
| mPC4 x Pollination | | | | 0.691 | 0.406 |
| mPC1 x Soil type | | | | 1.667 | 0.197 |
| mPC2 x Soil type | | | | 0.210 | 0.646 |
| mPC3 x Soil type | | | | 0.229 | 0.632 |
| mPC4 x Soil type | | | | 0.053 | 0.817 |
| mPC1 x Herbivory | | | | 0.061 | 0.805 |
| **mPC2 x Herbivory** | | | | **7.995** | **0.005** |
| mPC3 x Herbivory | | | | 1.872 | 0.171 |
| mPC4 x Herbivory | | | | 0.047 | 0.828 |
| mPC1 x Pollination x Herbivory | | | | 0.043 | 0.835 |
| mPC2 x Pollination x Herbivory | | | | 2.049 | 0.152 |
| mPC3 x Pollination x Herbivory | | | | 2.368 | 0.124 |
| mPC4 x Pollination x Herbivory | | | | 2.684 | 0.101 |
| mPC1 x Pollination x Soil Type | | | | 1.114 | 0.291 |
| mPC2 x Pollination x Soil Type | | | | 0.186 | 0.667 |
| mPC3 x Pollination x Soil Type | | | | 0.471 | 0.429 |
| mPC4 x Pollination x Soil Type | | | | 0.106 | 0.745 |
| mPC1 x Soil Type x Herbivory | | | | 0.047 | 0.829 |
| mPC2 x Soil Type x Herbivory | | | | 0.029 | 0.865 |
| **mPC3 x Soil Type x Herbivory** | | | | **5.484** | **0.019** |
| mPC4 x Soil Type x Herbivory | | | | 0.748 | 0.387 |
| mPC1 x Pollination x Soil Type x Herbivory | | | | 2.477 | 0.115 |
| mPC2 x Pollination x Soil Type x Herbivory | | | | 0.013 | 0.911 |
| **mPC3 x Pollination x Soil Type x Herbivory** | | | | **6.893** | **0.009** |
| mPC4 x Pollination x Soil Type x Herbivory | | | | 2.132 | 0.144 |
